# Hydrogen sulfide promotes nodulation and nitrogen fixation in soybean-rhizobia symbiotic system

**DOI:** 10.1101/367896

**Authors:** Hang Zou, Ni-Na Zhang, Qing Pan, Jian-Hua Zhang, Juan Chen, Ge-Hong Wei

## Abstract

The rhizobium-legume symbiotic system is crucial for nitrogen cycle balance in agriculture. Hydrogen sulfide (H_2_S), a gaseous signaling molecule, may regulate various physiological processes in plants. However, whether H_2_S has regulatory effect in this symbiotic system remains unknown. Herein, we investigated the possible role of H_2_S in the symbiosis between soybean (*Glycine max*) and rhizobium (*Sinorhizobium fredii*). Our results demonstrated that exogenous H_2_S donor (sodium hydrosulfide, NaHS) treatment promoted soybean growth, nodulation and nitrogenase (Nase) activity. Western blotting analysis revealed that the abundance of nitrogenase component nifH was increased by NaHS treatment in nodules. Quantitative real-time PCR data showed that NaHS treatment up-regulated the expressions of symbiosis-related genes *nodC* and *nodD* of *S. fredii*. Besides, expression of soybean nodulation marker genes including early nodulin 40 (*GmENOD40*), ERF required for nodulation (*GmERN*), nodulation signaling pathway2b (*GmNSP2b*) and nodulation inception genes (*GmNIN1a, GmNIN2a* and *GmNIN2b*) were up-regulated. Moreover, the expressions of glutamate synthase (*GmGS*), nitrite reductase (*GmNiR*), ammonia transporter (*GmSAT1*), and *nifH* involved in nitrogen metabolism were up-regulated in NaHS-treated soybean roots and nodules. Together, our results suggested that H_2_S may act as a positive signaling molecule in soybean-rhizobia symbiotic system and enhance their nitrogen fixation ability.

**Highlight:** We demonstrated for the first time that H_2_S as a signaling molecule may promote the establishment of symbiotic relationship and nitrogen fixation ability in the soybean-rhizobia symbiotic system.

## Introduction

Symbiosis is ubiquitous in terrestrial, freshwater, and marine communities. It has played a key role in the emergence of major life forms on Earth, and in the generation of biological diversity (Moran, 2006). Among all plant-microorganism symbioses, mutualism between legumes and bacteria known as rhizobia, is the most well-studied (Caballero-Mellado & Martinez-Romero, 1999). Symbiotic infection by rhizobia into legume roots leads to the formation of a specialized organ known as root nodule in which rhizobia differentiate into nitrogen-fixing bacteroids (Mergaert et al., 2006). Root nodules not only create an optimized environment for nitrogen fixation, but also provide a site where substance exchange can take place in plants (Becana & Sprent, 1987). In exchange for atmospheric nitrogen fixed by bacteroids, legumes provide energy and carbon sources to the bacterial partner (Werner et al., 2015).

In recent years, overuse of nitrogen fertilizers aimed at increasing soybean (*Glycine max*) and other crop yields has led to widespread environmental problems (Tian et al., 2012). Counterproductively, nitrogen fertilizer overuse does not promote soybean production since the redundancy of inorganic nitrogen in the soil layer surrounding roots inhibits nodulation and biological nitrogen fixation (Tilman et al., 2002). This reduced the utilization efficiency of nitrogen fertilizers and caused acidification and hardening of soil, along with various environmental problems. Thus, enhancing the nitrogen-fixing ability of the soybean-rhizobia symbiotic system is a practical solution for reducing the overuse of nitrogen fertilizers and preventing subsequent environmental problems (Masclaux-Daubresse et al., 2010).

The establishment of symbiosis between plants and microorganisms involves a complex regulatory network that includes nitric oxide (NO) and reactive oxygen species (ROS) (Li et al., 2011; Fukuto et al., 2012; Oldroyd, 2013). NO is produced in functional root nodules and is crucial for the establishment of symbiosis between *Medicago truncatula* and *Sinorhizobium meliloti* (Baudouin et al., 2006; Del Giudice et al., 2011). The production of NO during the infection process indicates a role for this gaseous molecule in recognition between plants and their bacterial partners (Hichri et al., 2016). Similar findings reported by Pii et al. (2007) suggested that NO and auxins may control the formation of indeterminate nodules in *M. truncatula*. Meanwhile, NO is also crucial for the development of functional nodules in soybean, as demonstrated by Leach et al. (2010) using a NO synthase-specific inhibitor. Furthermore, ROS and NO may synergistically control the early stages of the formation of legume-rhizobia symbiosis (Damiani et al., 2016). Additionally, NO have a negative effect on nitrogenase (Nase) activity in plants (Meilhoc et al., 2010; Cam et al., 2012).

Recently, the role of hydrogen sulfide (H_2_S), as a signaling molecule, regulating physiological processes in plants has become a hot topic. In addition to participating in adventitious root and lateral root formation and seed germination (Zhang et al., 2008; Zhang et al., 2009; Lin et al., 2012), H_2_S is reportedly involved in alleviating oxidative damage caused by heavy metals such as aluminum, copper, boron, cadmium, and chromium (Zhang et al., 2008; Wang et al., 2010; Li et al., 2012; Chen et al., 2013; Sun et al., 2013; Fang et al., 2016). Moreover, H_2_S also acts as an antioxidant signaling molecule to modulate ROS and antioxidant levels, thereby improving drought tolerance in soybean (Zhang et al., 2010). Alternatively, H_2_S enhances the drought tolerance of plants by affecting the biosynthesis and expression levels of genes associated with polyamines and soluble sugars (Chen et al., 2016). H_2_S also appears to alleviate saline-induced stress responses in bermudagrass (*Cynodon dactylon*) (Shi et al., 2013). Shen et al. (2013) found that H_2_S conferred heat tolerance in *Arabidopsis thaliana* through affecting microRNA expression. Moreover, H_2_S regulates stomatal movements together with NO, abscisic acid and inward-rectifying K^+^ channels (García-Mata & Lamattina, 2010; Scuffi et al., 2014; Papanatsiou et al., 2015). Chen et al. (2011) reported that H_2_S promotes photosynthesis by increasing ribulose-1, 5-bisphosphate carboxylase activity, and through thiol redox modification in *Spinacia oleracea*. Other studies indicate a role for H_2_S in autophagy (Álvarez et al., 2012; Romero et al., 2014; Laureano-Marín et al., 2016). In both animals and plants, most H_2_S responses are related to, or mediated by, NO or ROS (Zhang et al., 2009; Li et al., 2013; Scuffi et al., 2014; Laureano-Marín et al., 2016). However, whether H_2_S exerts synergistic or similar functions in the formation of legume-rhizobia symbiosis remains unclear.

In the present study, we found that treatment with an H_2_S donor (sodium hydrosulfide, NaHS) promotes infection of *S. fredii* into soybean roots, and enhances the biological nitrogen fixation ability of the symbiotic system by up-regulating symbiosis and nitrogen fixation related genes and proteins. Stimulation of nitrogen fixation and metabolism led to enhanced nitrogen assimilation and photosynthesis in soybean plants, which eventually promoted plant growth. This is the first study to investigate the regulatory role of H_2_S in the legume-rhizobia symbiotic system, and our findings suggest H_2_S is a positive regulator in the establishment of symbiosis and symbiotic nitrogen fixation between *G. max* and *S. fredii*. This research will likely inspire future study of the role of H_2_S in plant physiology, and might offer a possible solution to increasing soybean production.

## Materials and methods

### Plant growth and H_2_S treatment

Soybean (*Glycine max* cv. Zhonghuang 13) seeds were surface-sterilized with 75% ethyl alcohol and sodium hypochlorite, then placed on a 1% agar plate for 72 h at 28°C in the dark. 800 ml of growth medium (vermiculite and perlite, v:v = 1:1) was watered with 400 ml nitrogen-free nutrient solution (100 mg/L CaCl_2_, 100 mg/L KH_2_PO_4_, 50 mg/L Ferric citrate, 150 mg/L NaH_2_PO_4_, 120 mg/L MgSO_4_·7H_2_O, 2.86 mg/L H_3_BO_3_, 2.3 mg/L MnSO_4_·4H_2_O, 2.8mg/L ZnSO_4_·7H_2_O, 13 mg/L Na_2_MoO_4_·2H_2_O, 2.2 mg/L CaSO_4_·5H_2_O) and sterilized in a polypropylene planting bag. Germinated seeds were transferred into growth medium (one seeding/bag). Seven-day-old soybean seedlings were divided into four groups: first group served as controls (CK), second group was treated with 100 μM NaHS (H), third group was inoculated with rhizobia (*Sinorhizobium fredii* Q8 strain) (Q), fourth group inoculated with *S. fredii* and treated with NaHS (QH). All seedlings in the Q and QH groups were inoculated with 10 mL of a rhizobial suspension (OD_600_ = 0.5) when the first main leaves of plants were fully expanded. NaHS was used as the H_2_S donor (Christou et al., 2013). Seedlings in the H and QH groups were watered with 10 mL of NaHS solution (100 μM) every 3 days until harvest, and seedlings from the other two groups were watered with double distilled H_2_O instead. 50 ml of sterile nitrogen-free nutrient solution was added into each bag every 7 days to maintain the steady humidity and ionic concentration. Twenty seedlings were harvested every 7 days since inoculation from each group, and half of the samples were dried to a constant weight for dry matter determination, while the other half was immediately frozen in liquid nitrogen and stored at −80°C.

### Root length, shoot length, root weight, and shoot weight measurements

The longest distance of root/shoot tip to the junction of root and shoot were set as root length and shoot length. Twenty soybean seedlings of each treatment group were used as replicates for these measurements.

### Nitrogenase activity determination

Nitrogenase (Nase) activity was quantified using the acetylene reduction method by Fishbeck et al. (1973) with slight modifications. Fresh soybean root nodules were transferred into a 10 mL rubber-capped airtight glass bottle filled with a mixture of acetylene and air (v:v = 1:100). Bottles were incubated at 28°C for 3 h, and the content of ethylene was determined using a gas chromatography system (Agilent Technologies, La Jolla, CA, USA).

### Infection event assay

The *S. fredii* Q8 strain harboring the enhanced green fluorescence protein encoded on the pMP2444 plasmid was employed. The plasmid was incorporated into the Q8 strain by tri-parent hybridization through the assistant plasmid pRK2013 (Nishikawa et al., 2008). Soybean roots were inoculated with a rhizobial suspension as described above, and typical infection events were determined at 5 and 7 DPI using a BX53 fluorescence microscope (Olympus, Tokyo, Japan).

### Development of a fluorescence probe for H_2_S application and fluorescent intensity quantification

H_2_S fluorescent probe SF7-AM was purchased from Sigma-Aldrich (CAS: 1416872–50-8; Dallas, TX, USA). As described by Lin et al. (2013), plant tissues were incubated in 5 mM SF7-AM for 1 h, washed with 20 mM HEPES, and visualized and photographed using a BX53 fluorescence microscope (Olympus, Tokyo, Japan). The fluorescence intensity was quantified by ImageJ software (National Institutes of Health, Bethesda, MD, USA).

### Electron microscopy observation of nodule micromorphology

Root nodules observed using a transmission electron microscope (FEI, Czech, USA). Plant tissues treated with/without 100 μM NaHS were gently washed, and clean root nodules were cut into tiny slices, prefixed in 4% glutaraldehyde, washed with 0.1 M pH 6.8 phosphate buffer, and post-fixed in 1.0% osmium tetroxide. After at least 3 h of fixation, nodules were dehydrated in an ascending ethanol series (Yuan et al. 2017), and embedded in LR white resin. Finally, thin sections were excised from the embedded samples using an ultramicrotome equipped with a glass knife, and ultrathin sections were mounted on copper grids for transmission electron microscopy examination.

### Chlorophyll content determination

Measurement of the chlorophyll content in soybean seedlings was carried out using a SPAD-502 Plus leaf chlorophyll meter (Konica Minolta, Kumamoto, Japan). A total of 20 leaves (third fully developed leaf) from 10 soybean seedlings per treatment were measured at 10:00 am every 7 days until harvest.

### Determination of photosynthesis and chlorophyll fluorescence parameters

Photosynthetic rate (Pn), stomatal conductance (Gs), carbon dioxide concentration (Ci), transpiration rate (Tr) and vapor pressure divicit (Vpdl) were measured using a portable photosynthesis system (Li-6400, LiCor, Lincoln, NE, USA) on the second fully expanded leaf of soybean seedling. Ten different leaves from ten different soybean seedlings were set as replicates, and the measurement were conducted three times. Air temperature, light intensity, CO_2_ concentration, and air relative humidity were maintained at 25°C, 800 μmol m^−2^ s^−1^ PAR, 380 μL l^−1^, and 90%, respectively. Determination was conducted from 9:00 am to 11:00 am to avoid high temperature and air vapor pressure deficits. Light was supplemented using an LED light system. Vapor pressure deficit during measurement was ∼1 kPa.

Chlorophyll fluorescence measurement was conducted on 10 leaves (third fully developed leaf) from 10 soybean seedlings using a Plant Efficiency Analyzer (Hansatech Instruments Ltd., Norfolk, England). Before measurement, leaves were pretreated in the dark for 30 min. Fo, Fm, Fv (= Fm - Fo), and Fv/Fm parameters were recorded for 15 s at a photon flux density of 4000 μmol m^−2^ s^−1^. Additionally, we measured the steady-state fluorescence level (Fs′) under continuous illumination, the maximal fluorescence level (Fm′) induced by a saturating light pulse at the steady-state, and the minimum fluorescence level (Fo′) after exposure to far-red light for 3 s. Then, Fv′ was calculated using the formula Fv′ = Fm′ - Fo′. PSII was calculated using the formula PSII = 1 - (Fs′ / Fm′). The ETR was calculated using the formula ETR = PAR × PSII × 0.85 × 0.5. NPQ was calculated using the formula NPQ = (Fm / Fm′) - 1. qP was calculated using the formula qP = (Fm′ - Fs) / (Fm′ - Fo′) (Krause & Weis, 2003).

### Sodium dodecyl sulfate-polyacrylamide gel electrophoresis and western blotting assays

Total protein of soybean samples was extracted following the protocol of Chen et al. (2011). After total protein isolation, protein concentrations were quantified using the Bradford (1976).

For western blotting analysis, proteins (200 μg for each sample) were separated by SDS-polyacrylamide gel electrophoresis (PAGE) using 12% acrylamide gels, and transferred to a polyvinylidene difluoride membrane. The membrane was blocked overnight with 5% skim milk powder. Protein blots were probed with primary antibodies for nifH (1:2000, Agrisera, Sweeden, AS01 021A) or CHS (1:1000, Agrisera, AS12 2615) at a dilution of 1:5000 or 1:3000, respectively, at 4°C overnight. Blots were washed three times in TBST solution (50 mM TRIS-HCl pH 8.0, 150 mM NaCl, 0.05% tween-20), followed by incubation with secondary antibody (anti-rabbit IgG horse radish peroxidase-conjugated, Sungene, China, 1:5000 dilution) overnight at 4°C. Actin (1:5000; Agrisera, AS13 2640) was used as an internal control. Blots were finally washed with PBST three times, and imaged using a Molecular Imager Gel Doc XR System (BioRad, Hercules, CA, USA). The optical density value was determined using Image J software and used to estimate the protein abundance.

### Total RNA isolation, reverse transcription, and gene expression analysis

Total RNA isolation was performed using the TaKaRa MiniBEST Plant RNA Extraction Kit (Takara, Dalian, China) according to the manufacturer’s instructions. RNA integrity was examined by 1% agarose gel electrophoresis. RNA concentration was determined using an Epoch Microplate Spectrophotometer (BioTek, Winooski, VT, USA). Reverse transcription was conducted using TaKaRa PrimerScript TM RT master Mix (Takara) as suggested by the manufacturer. qRT-PCR was carried out with a Quantstudio 6 Flex real-time PCR system (Thermo Fisher, Carlsbad, CA, USA) and SYBR Premix Ex Taq II (Takara). The qPCR program are described in Table S2. Primer sequences are listed in Table. S1.

### Statistical analysis

Statistical significance was tested by analysis of variance using SPSS 19.0 (SPSS Inc., Chicago, IL, USA). All the significances in time course experiments were tested using the repeated measured ANOVA of general linear model, for these parametric tests, differences were considered statistically significant at *P* < 0.05.

## Results

### Exogenous addition of H_2_S donor promotes soybean plants growth

To explore the regulatory effects of H_2_S on soybean plant growth and symbiosis with *S. fredii*, 10 mL of 100 μM NaHS solution was used to treat each soybean roots, and the growth of roots and shoots was measured after inoculation. In both inoculated and non-inoculated groups, NaHS treatment promoted root elongation (Fig. 1A). In the inoculated plants, average root length was elevated by H_2_S treatment at most of the checkpoints, and this increase in root length was turned out to be statistically significant (*F* = 4.876, *P* = 0.046) by repeated measured (RM) ANOVA of general linear model (GLM). In the non-inoculated plants, NaHS treatment also slightly extended the average root length, but the extension of root length by NaHS treatment was not significant (RM GLM, *F* = 3.183, *P* = 0.087). Similar results were observed in soybean shoots. Shoot length curve indicated that NaHS treatment did not have a significant effect on shoot elongation in non-inoculated soybean plants (RM GLM, *F* = 2.061, *P* = 0.193). However, in the inoculated soybean plants, the average shoot length was significantly increased by NaHS treatment (*F* = 4.294, *P* = 0.048) (Fig. 1B). The biomass of dry matters in inoculated soybean roots was significantly higher compared with non-treated plants (RM GLM, *F* = 12.323, *P* = 0.013). However, there was no significant difference observed among the other three treatments (Fig. 1C). NaHS treatment only provided a very slight and not significant promotion in shoot dry weight compared with respective controls in both inoculated and non-inoculated plants (RM GLM, *F* = 0.973, *P* = 0.392; *F* = 2.986, *P* = 0.089) (Fig. 1D). Taken together, these results demonstrated that NaHS treatment promoted the growth of soybean seedlings, especially the growth of soybean roots under symbiotic conditions.

**Figure 1.**
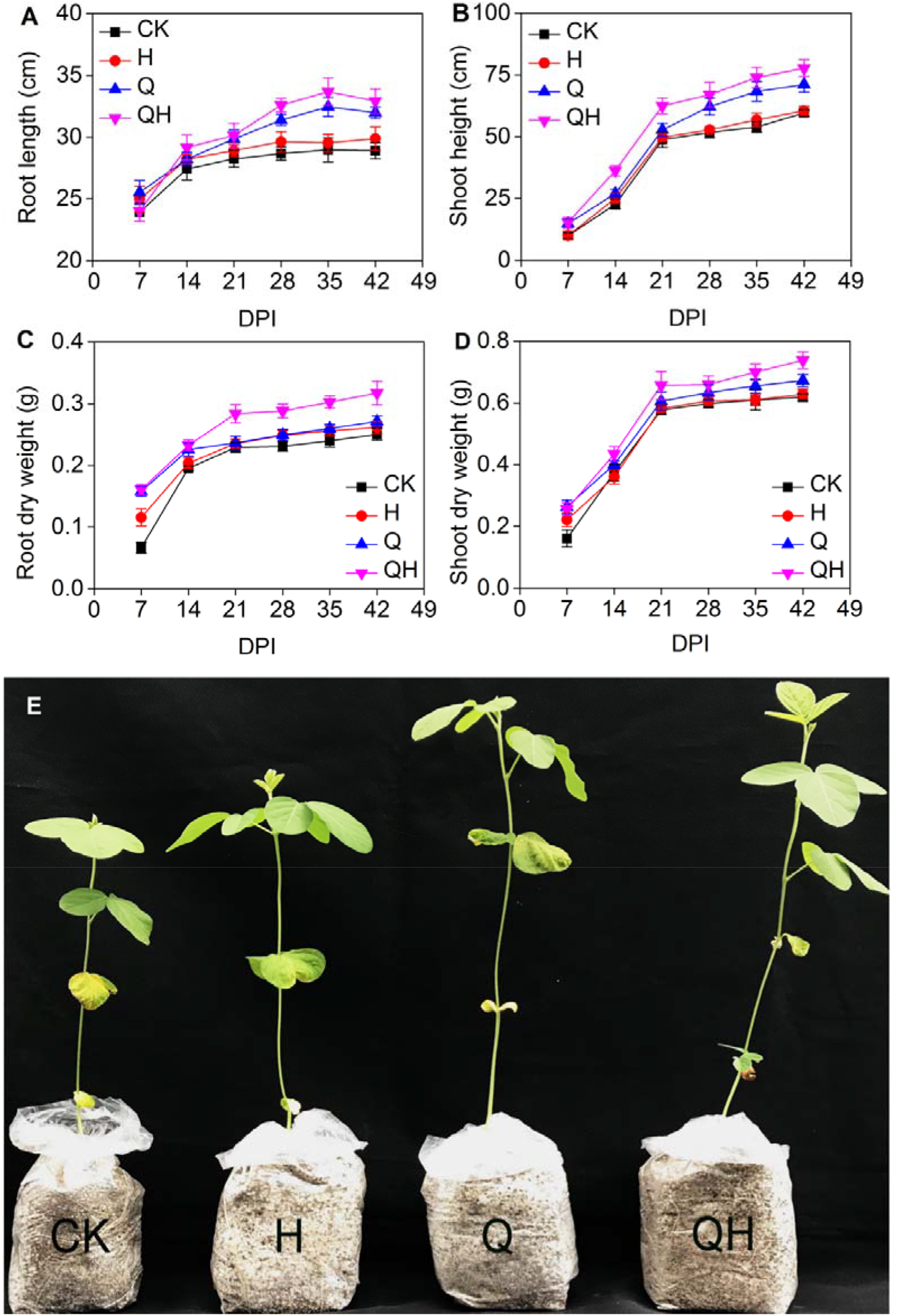
H_2_S’s effect on the growth of soybean seedlings. Time course of root and shoot length (A, B) and dry weight (C, D) for soybean plants. (E) Soybean plants following different treatments at 14 days post-inoculation (DPI). Soybean plants were harvested every 7 days since inoculation. Values are means ± SE (n = 20). CK, Controls. H, 100 μM NaHS. Q, Soybean seedlings inoculated with *Sinorhizobium fredii* Q8 strain. QH, Soybean seedlings inoculated with *S. fredii* Q8 strain and treated with 100 μM NaHS.

### Exogenous addition of the H_2_S donor promotes rhizobial infection, root nodulation, and nitrogenase activity

We counted the number of nodules on each harvested soybean root at 7, 14, 21, 28, 35 and 42 DPI to determine the effects of NaHS on nodulation in soybean plant. The result showed that H_2_S promoted soybean nodulation (RM GLM, *F* = 9.74, *P* = 0.019, Fig. 2A). Meanwhile, the addition of NaHS promoted not only soybean nodulation, but also the N fixation ability of these nodules. Nase activity assay which was measured by the acetylene reduction (AR) experiments showed that NaHS treatment significantly increased Nase activity in root nodules (RM GLM, *F* = 22.874, *P* = 0.007) (Fig. 2B). The AR ability showed significant increase compared with the control, these results suggested that H_2_S stimulated soybean nodulation and enhanced the N fixing potential of the soybean-rhizobia symbiotic system.

**Figure 2.**
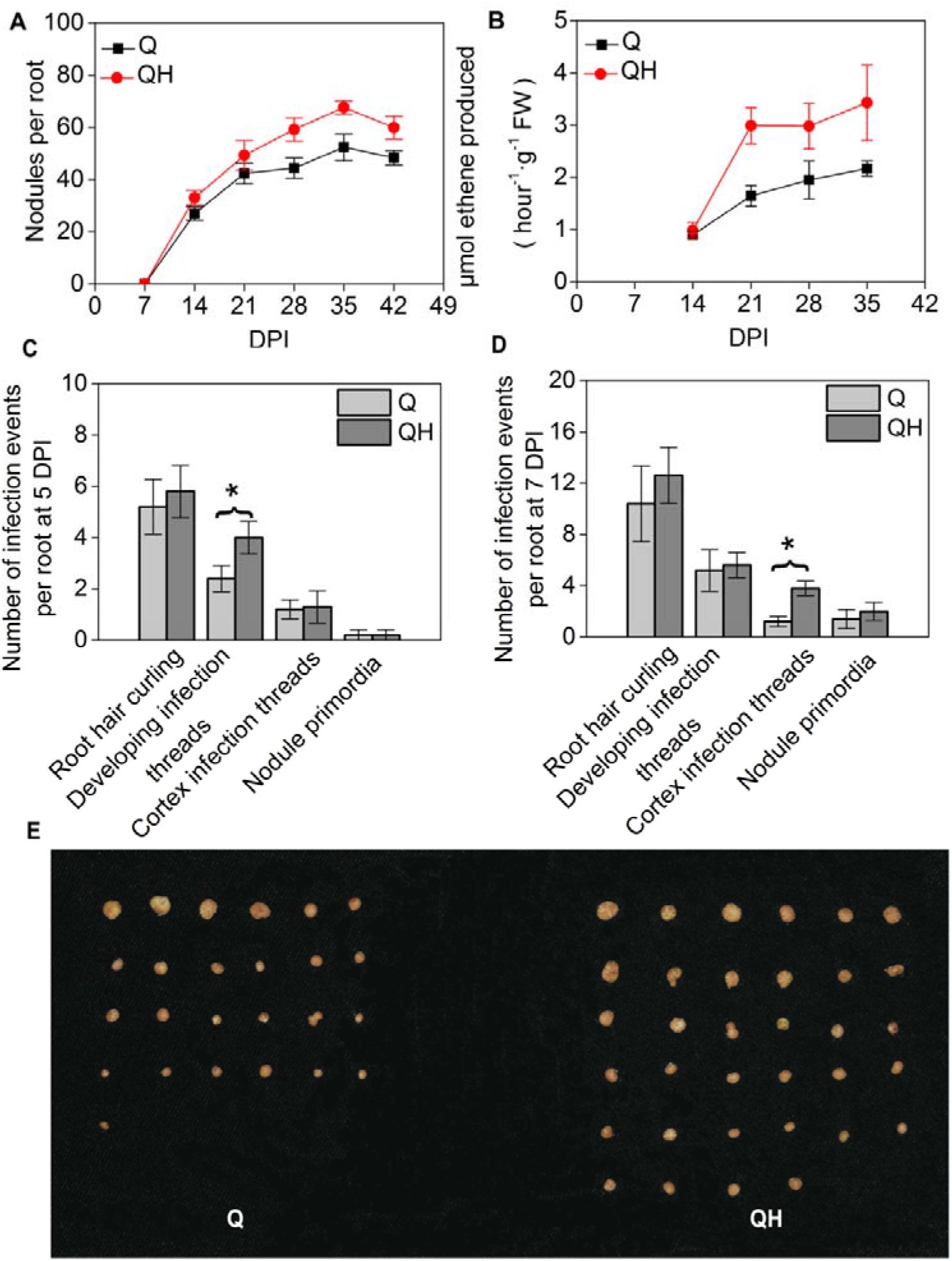
H_2_S’s effect on soybean nodulation and nitogenase (Nase) activities. Root nodule amount (A) and nitrogenase activity assay (B). Infection events assay at 5 DPI (C) and 7 DPI (D). (E) indicates red functional nodules obtained from soybean roots at 14 DPI. For nodule amount, values are means±SE (n=20). For nitrogenase activity, values=means ± SE (n=5). * *p* < 0.05. Q: Soybean seedlings inoculated with *Sinorhizobium fredii* Q8 strain. QH: Soybean seedlings inoculated with *S. fredii* Q8 strain and treated with 100 μM NaHS.

Additionally, another phenotypic analysis of infection events was conducted to further delineate the effect of NaHS treatment on soybean nodulation. Root hair curling, infection threads, and nodule primordia formed during the nodulation process were observed and counted (Fig. 2C and Fig. S4). At 5 DPI, more root hair curling and developing infection threads were found in NaHS-treated soybean roots (Fig. 2C). However, no significant difference in number of cortex infection threads and nodule primordia were found between NaHS-treated and untreated soybean roots. At 7 DPI, cortex infection threads in NaHS-treated soybean roots were more abundant than those in untreated controls (Fig. 2D). In addition to enhanced nodulation in NaHS-treated soybean plants, the average size of red mature root nodules on soybean roots harvested at 14 DPI was noticeably larger than on untreated control roots (Fig. 2E). These results suggested that H_2_S may act as a positive regulator of soybean nodulation.

### Exogenous addition of an H_2_S donor increases endogenous H_2_S concentration in soybean roots and nodules

To confirm that the observed changes were caused by the H_2_S released from NaHS, we used an H_2_S-specific fluorescence probe to measure the amount of endogenous H_2_S in soybean plant tissues including lateral root, main root, and nodule. Endogenous H_2_S was measured in different soybean tissues after loading with the sulfidefluor-7-acetoxymethyl ester (SF7-AM) probe (Fig. 3 and Fig. S3). Before NaHS treatment, the observed low fluorescence intensity indicated a low concentration of endogenous H_2_S in these tissues (Fig. 3G-I). After treatment with 100 μM NaHS, higher fluorescence intensity was observed, indicating that NaHS significantly increased the endogenous H_2_S concentration in these tissues (Fig. 3J-L). Due to the drastic increment of fluorescent intensity by the NaHS treatment, we may rule out the possibility of this strong fluorescent was caused by autofluorescent. The effect of NaHS treatment on endogenous H_2_S concentration in plant tissues was also verified by quantification of the fluorescence intensity in these tissues (Fig. S3). Together, these results indicated that NaHS treatment significantly increased the endogenous H_2_S concentration in soybean tissues.

**Figure 3.**
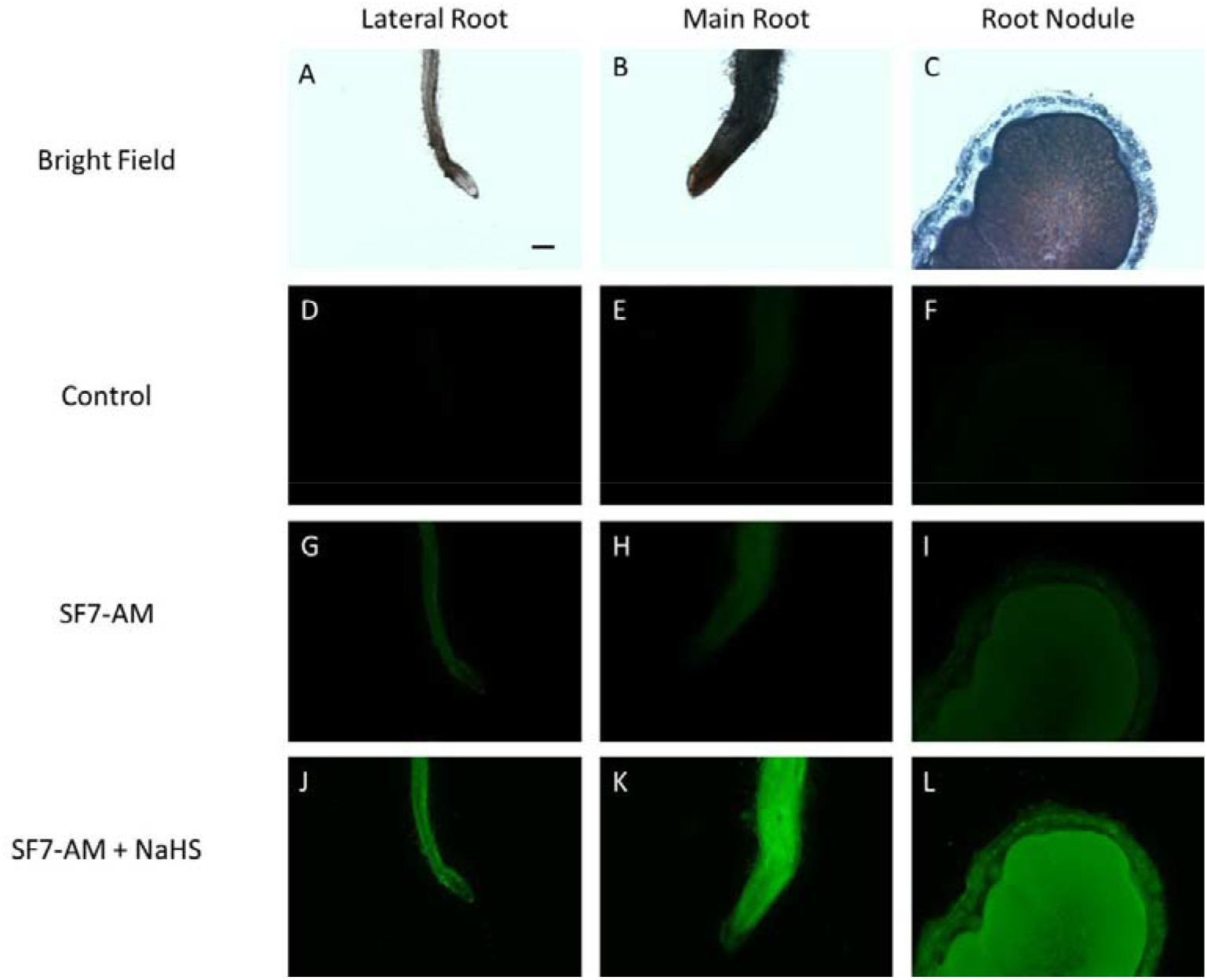
Fluorescence probe assay of endogenous H_2_S in different soybean plant tissues. (A–C), bright field images of untreated plant tissues. (D–F), fluorescence field images of untreated plant tissues. (G–I), Fluorescence field images of tissues loaded with H_2_S-specific fluorescence probe SF7-AM. (J–L), Tissues loaded with SF7-AM and treated with 100 μM NaHS. Bar = 200 μm.

### H_2_S influenced bacteroids colonization

We used light microscope to determine whether H_2_S could lead to structural changes in nodules. However, the paraffin sections of nodules obtained at different time did not exhibit any significantly difference between NaHS treated nodules and control nodules (Fig. S5).

Transmission electron microscopy was then conducted to observe the microscopic structure of infected cell bacteroids in nodules. Bacteroids in nodules harvested at 7, 21, and 28 DPI exhibited typical morphological characteristics (data not shown), and no obvious differences were found between NaHS-treated nodules and controls. However, at 14 DPI, intercellular colonization of bacteroids was observed in NaHS-treated nodules (Fig. 4B, indicated by dark triangles). Typical bacteroids were present inside nodule cells and surrounded by a plant-derived peribacteroid membrane (indicated by a dark arrows in Fig. 4A, B), but separated bacteroids in the intercellular space lacked the typical peribacteroid membrane structure compared with the control nodules harvested at 14 DPI (Fig. 4A). These results implied that H_2_S may be involved in the regulation of rhizobial differentiation in soybean nodules.

**Figure 4.**
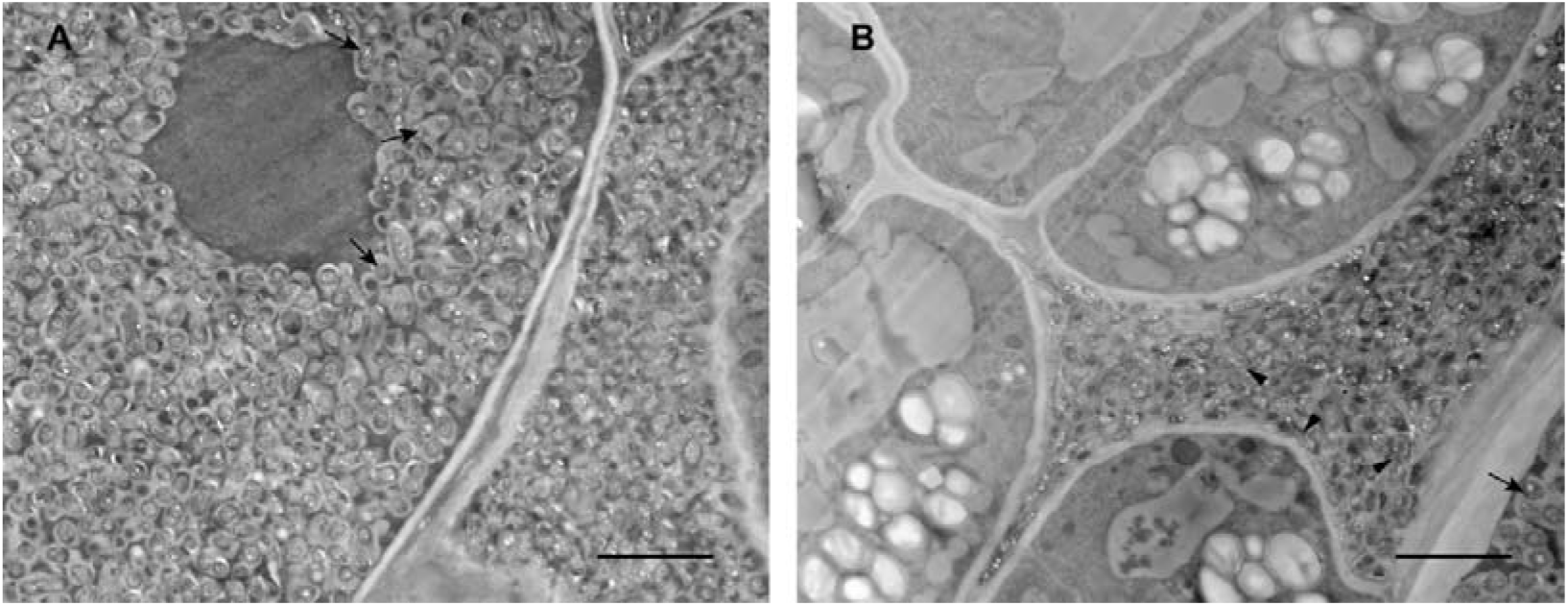
Transmission electron microscopy analysis of nodule ultrastructure. Nodule of control soybean seedling (A). Nodule of soybean seedling treated with100 μM NaHS (B). Nodules were obtained at 14 days post-inoculation. Black arrows indicate bacteroids in infected cells, and black triangles indicate intercellular bacteroids. Bars = 5 μm.

### Chlorophyll content, photosynthesis intensity, and photosystem II activity are elevated by H_2_S

NaHS treatment increased the chlorophyll content in both inoculated and non-inoculated plants from 7 to 42 DPI. In the inoculated soybean plants, the chlorophyll content in soybean leaves was significantly increased by NaHS treatment (RM GLM, *F* = 32.338, *P* = 0.004). While in the non-inoculated plants, NaHS did not make any significant difference (RM GLM, *F* = 2.004, *P* = 0.182) (Fig. 5A). Net photosynthetic rate (Pn) curve indicated that NaHS treatment enhanced the photosynthesis in soybean leaves in the inoculated plants, (RM GLM, *F* = 20.841, *P* = 0.023). Again, NaHS treatment did not influence Pn in non-inoculated plants (RM GLM, *F* = 2.332, *P* = 0.079) (Fig. 5B). Stomatal conductance (Gs) of both inoculated and non-inoculated plants was not affected by NaHS (RM GLM, *F* = 2.081, *P* = 0.164; *F* = 2.827, *P* = 0.095) (Fig. 5C). Furthermore, no significant difference of the intercellular carbon dioxide concentration (Ci) was observed in neither inoculated nor non-inoculated soybean plants (RM GLM, *F* = 2.339, *P* = 0.138; *F* = 1.887, *P* = 0.295) (Fig. 5D). Transpiration rate (Tr) and vapor pressure deficit of leaf (Vpdl) were not markedly regulated by NaHS treatment (Fig. 5E-F).

**Figure 5.**
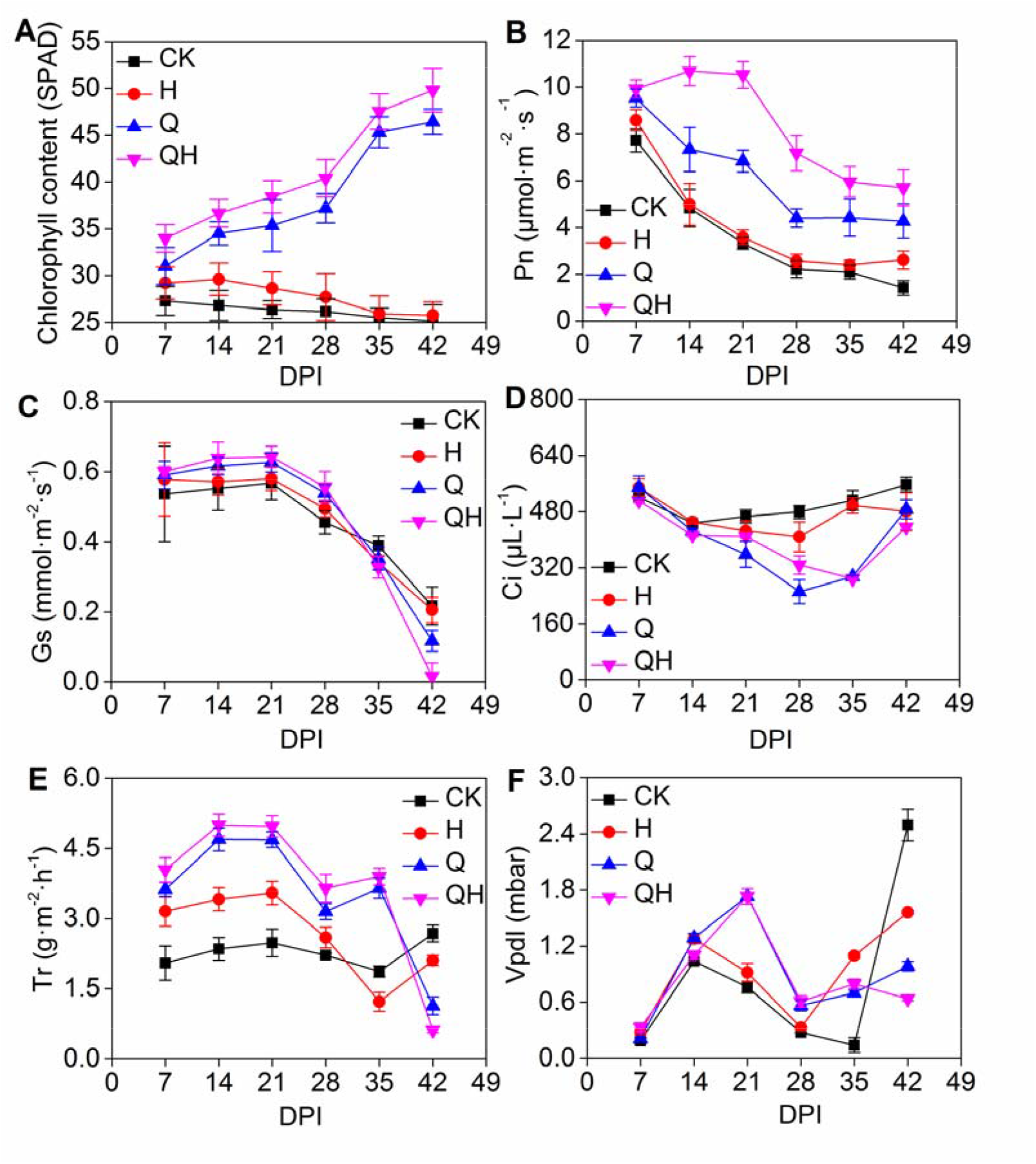
H_2_S’s effect on photosynthetic parameters. Chlorophyll content (A), net photosynthetic rate (Pn, B), stomatal conductance (Gs, C), carbon dioxide concentration (Ci, D), transpiration rate (Tr, E) and Vapor pressure divicit (Vpdl, F). Values are means ± SE (n = 10). CK, Controls. H, 100 μM NaHS. Q, Soybean seedlings inoculated with *Sinorhizobium fredii* Q8 strain. QH, Soybean seedlings inoculated with *S. fredii* Q8 strain and treated with 100 μM NaHS.

In terms of chlorophyll fluorescence parameters, the quantum yield of PSII photochemistry (PSII), the ratio of variable fluorescence to maximum fluorescence (Fv/Fm), and photochemical efficiency of PSII in the light (Fv′/Fm′) followed a similar trend after treatment with 100 μM NaHS (Fig. 6A, C, D). In the non-inoculated group, PSII in NaHS-treated plants was higher than controls at all checkpoints (RM GLM, *F* = 8.828, *P* = 0.036). Similarly, in the inoculated group, PSII in NaHS-treated plants was higher than controls (RM GLM, *F* = 5.352, *P* = 0.047) (Fig. 6A). Electronic transport ratio in inoculated plants was increased in the early stage of the NaHS treatment, while in the late stage, ETR was lower in NaHS-treated plants than control plants, however, no statistically significant difference was made by NaHS treatment (RM GLM, *F* = 0.963, *P* = 0.759). In non-inoculated plants, the effect of NaHS on ETR was not obvious (RM GLM, *F* = 1.023, *P* = 0.702) (Fig. 6B). Fv/Fm in inoculated plants was higher than in non-inoculated plants. However, in both inoculated and non-inoculated groups, there was not significantly difference between NaHS-treated and control plants during the entire testing period (RM GLM, *F* = 3.392, *P* = 0.055; *F* = 1.977, *P* = 0.183) (Fig. 6C). In non-inoculated plants, NaHS treatment did not affect Fv′/Fm′ (RM GLM, *F* = 1.339, *P* = 0.217). However, in inoculated plants, Fv′/Fm′ in NaHS-treated plants was higher than controls from (RM GLM, *F* = 9.521, *P* = 0.029) (Fig. 6D). NaHS treatment did not have a significant regulatory effect on non-photochemical quenching (NPQ) and photochemical quenching (qP) parameters in soybean plants (Fig. 6E, F).

**Figure 6.**
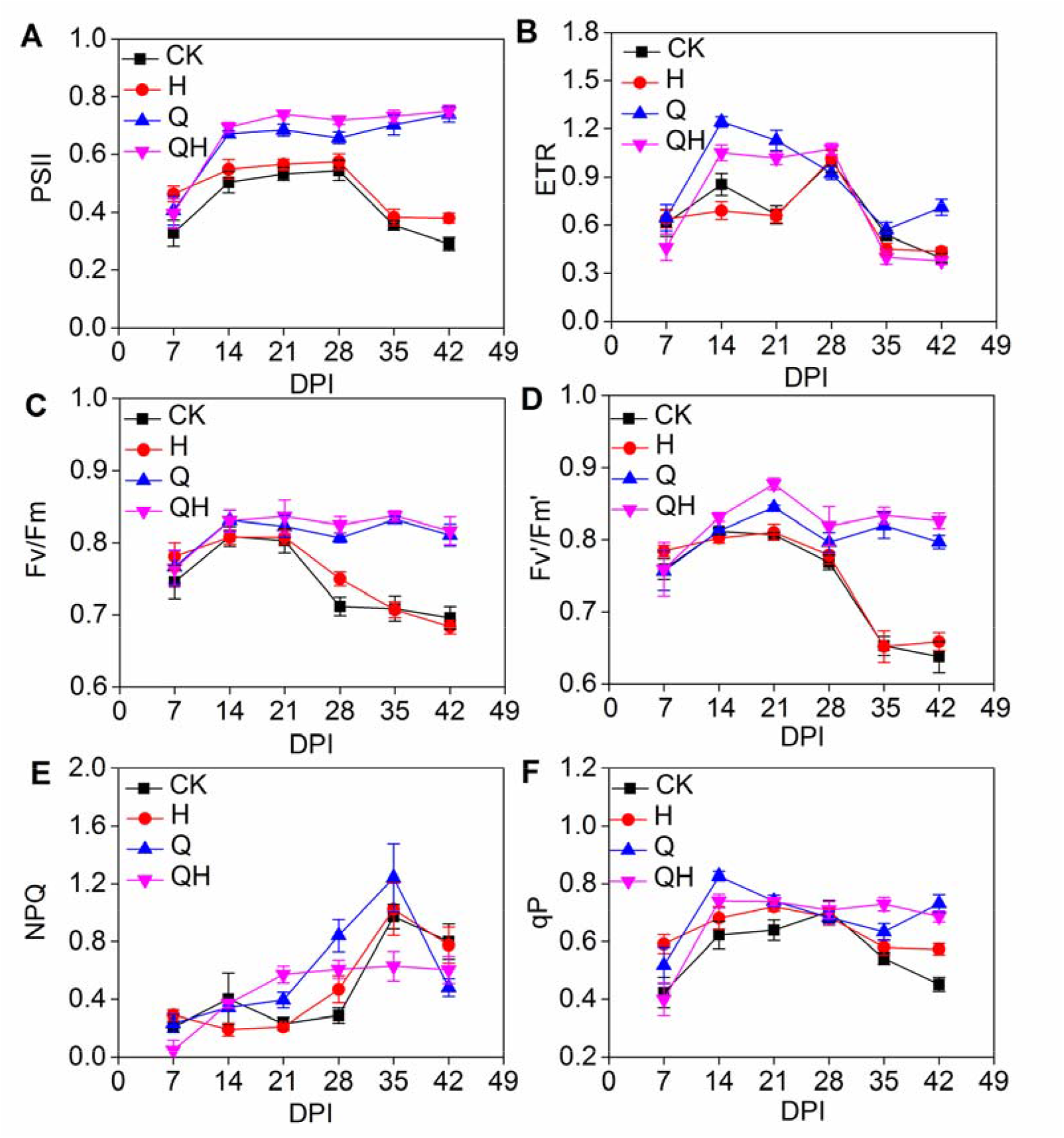
H_2_S’s effect on chlorophyll fluorescent parameters. Quantum yield of PSII photochemistry (PSII, A), electronic transport ratio (ETR, B), the ratio of variable fluorescence to maximum fluorescence (Fv/Fm, C), photochemical efficiency of PSⅡ in the light (Fv′/Fm′) (D), NPQ (E) and photochemical quenching (qP, F). Values are means ± SE (n = 10). CK, Controls. H, 100 μM NaHS. Q, Soybean seedlings inoculated with *Sinorhizobium fredii* Q8 strain. QH, Soybean seedlings inoculated with *Sinorhizobium fredii* Q8 strain and treated with 100 μM NaHS.

### H_2_S affects symbiosis and nirtogen fixation-related protein expression

Western blotting analysis of chalcone synthase (CHS) and the Nase iron-containing protein (nifH) were conducted to investigate the regulatory effects of H_2_S on the expression of key proteins involved in symbiotic establishment and nitrogen fixation. The result of western blotting indicated that H_2_S did not trigger any significant changes in the protein expression abundance of CHS in NaHS treated soybean root nodules compared with the non-treated controls (Fig. 7A, B). However, NaHS treatment significantly increased the protein expression level of nifH in root nodules. At 14, 21, 28, 35 DPI, the abundances of nifH protein were significantly higher than that in the control nodules (Fig. 7A, C).

**Figure 7.**
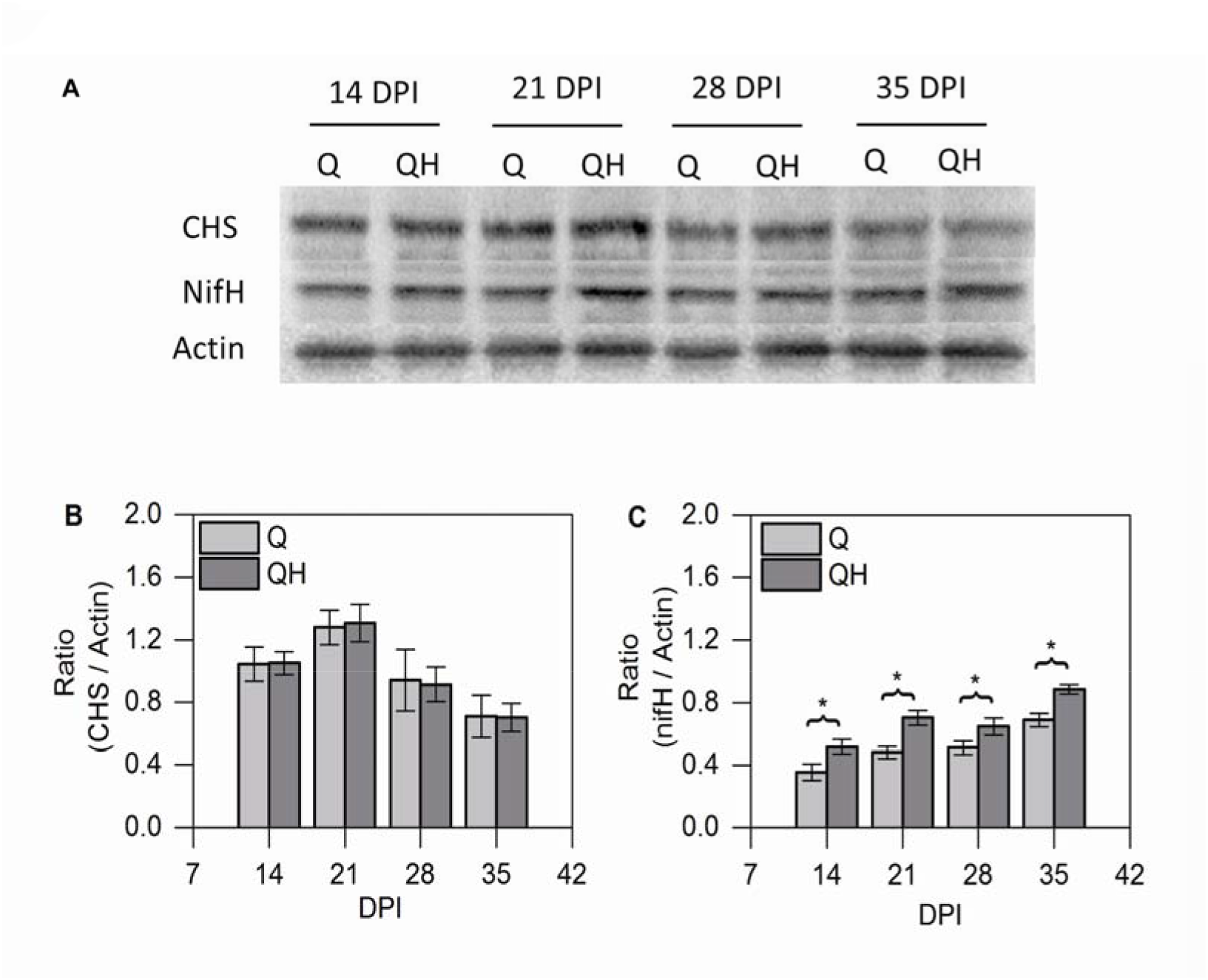
Western blotting analysis of the expression of chalcone synthase (CHS) and the nitrogenase iron-containing protein (nifH) in soybean nodules at 14, 21, 28, and 35 days post-inoculation (DPI; A). Expression levels of CHS and nifH are relative to actin (B and C). Values are means ± SE (n = 3). * *p* < 0.05. Q, Soybean seedlings inoculated with *Sinorhizobium fredii* Q8 strain. QH, Soybean seedlings inoculated with *Sinorhizobium fredii* Q8 strain and treated with 100 μM NaHS.

### RT-qPCR analysis of the expression of symbiosis-related gene

To further elucidate the mechanism that H_2_S promoted nodulation in soybean, we examined the expression profile of several nodulation marker genes in soybean roots including *GmENOD40, GmERN, GmNSP2b, GmNIN1a, GmNIN2a*, and *GmNIN2b*. We selected 12 h, 1 d, 3 d, 5 d and 7 d post inoculation of *S. fredii* as checkpoints. The qRT-PCR results demonstrated that H_2_S stimulated the expression levels of these genes in soybean roots. For instance, repeated measured ANOVA of general linear model suggested that in the inoculated soybean roots, NaHS treatment significantly elevated the expression levels of *GmENOD40* (*F*= 10.253, *P* = 0.027) (Fig. 8A) and *GmNIN1a* (*F* = 9.354, *P* = 0.039) (Fig. 8D) in the inoculated roots during the entire treatment period. As for *GmERN* and *GmNSP2b*, NaHS treatment led to significant up-regulation in their expression levels only from 12 h to 3 d post inoculation (*F* = 10.265, *P* = 0.007; *F* = 21.236, *P* = 0.01) (Fig. 8B, C). On the other hand, *GmNIN2a* and *GmNIN2b* expression in inoculated roots were stimulated by NaHS treatment only from 3 DPI to 7 DPI (*F* = 15.581, *P* = 0.017; *F* = 18.242, *P* = 0.013) (Fig. 8E, F). NaHS treatment did not give rise to any significant difference in the non-inoculated soybean roots.

**Figure 8.**
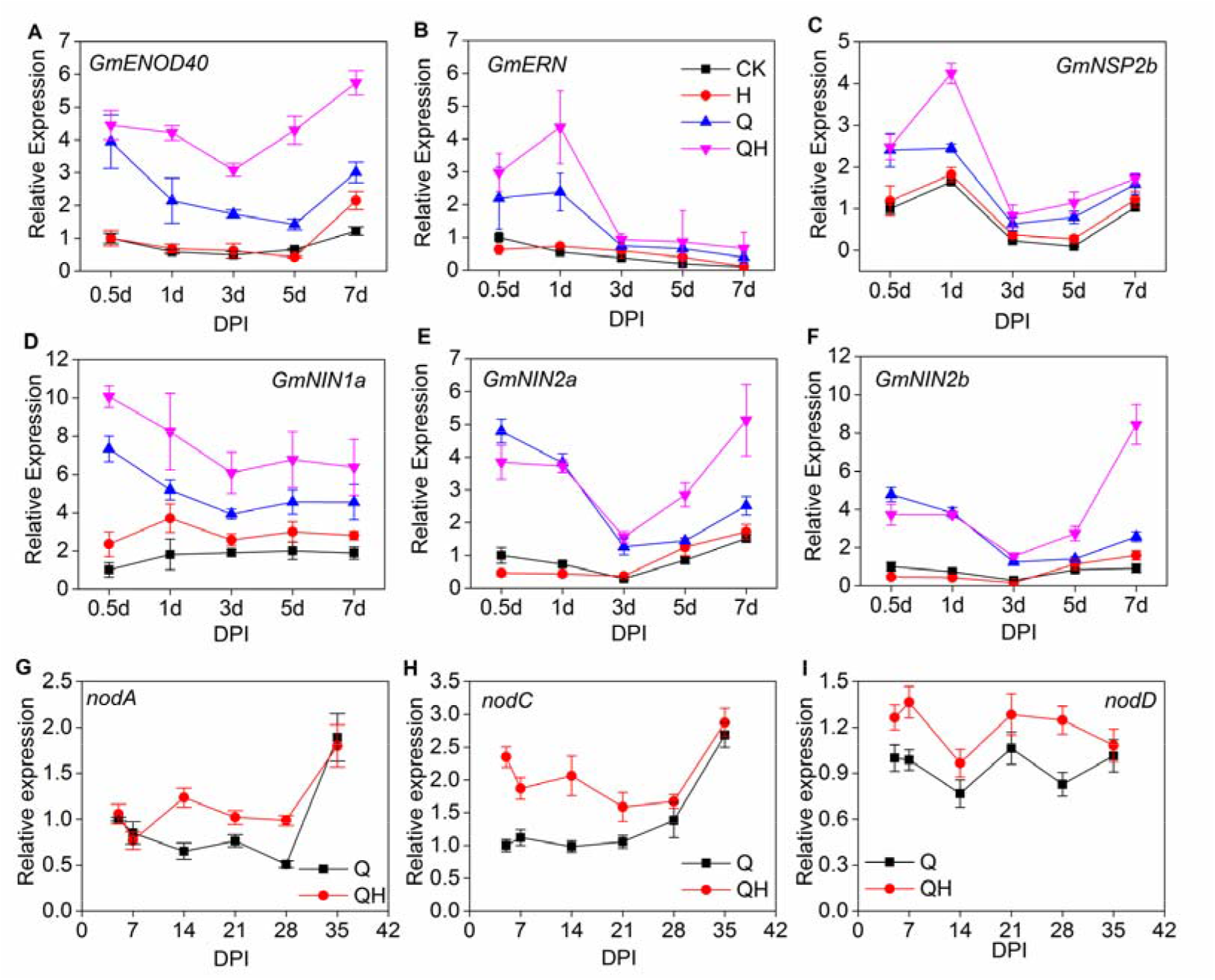
Gene expression level of symbiotic related genes. Relative expression levels of *GmENOD40* gene (A), *GmERN* gene (B), *GmNSP2b* gene (C), *GmNIN1a* gene (D), *GmNIN2a* gene (E), *GmNIN2b* gene (F), *NodA* gene (G), *NodC* gene (H) and *NodD* (I) gene are displayed in multiple line charts with symbols. CK, controls. H, soybean plants treated with 100 μM NaHS. Q, Soybean plants inoculated with the *Sinorhizobium fredii* Q8 strain. QH, Soybean plants inoculated with the *Sinorhizobium fredii* Q8 strain and treated with 100 μM NaHS. The expression of *GmENOD40, GmERN, GmNSP2b* and *GmNIN* genes were relative to the expression level of respective gene in CK roots at 0.5 DPI. The expression of *nodA, nodC* and *nodD* were relative to the expression level of respective gene in Q nodules at 5 DPI. Values are means ± SE (n = 9).

The expression of *nodA, nodC* and *nodD* genes of *S. fredii* was examined to verify whether NaHS treatment could also induce the symbiotic reaction in rhizobia. In this quantitive assay, root nodules were used and 5, 7, 14, 21, 28 and 35 DPI was selected as checkpoints. qRT-PCR results suggested that the expression of *nodC* and *nodD* were significantly induced by NaHS treatment (RM GLM, *F* = 5.617, *P* = 0.036; *F* = 11.338, *P* = 0.021) (Fig. 8H, I). Though the relative expression levels of *nodA* was also up-regulated in the NaHS-treated soybean nodules, but the difference failed to exhibit statistical significances within the entire treatment period (RM GLM, F = 3.336, P = 0.069) (Fig. 8G).

### RT-qPCR analysis of the relative expression of key enzymes related to nitrogen metabolism

Expression of N-metabolism related genes was investigated to determine the possible influence of H_2_S on the molecular mechanisms of symbiotic nitrogen fixation and nitrogen metabolism in soybean plants. Glutamate synthase (*GmGOGAT*), asparagine synthase (*GmAS*), nitrite reductase (*GmNiR*), ammonia transporter *SAT1* (*GmSAT1*), leghemoglobin (*GmLb*) in soybean plant, and *nifH* in *S. fredii* were selected for qRT-PCR analysis.

The transcript abundances of *GmGOGAT* is higher during the entire NaHS treatment period than that in control plant (RM GLM, *F* = 7.136, *P* = 0.047) (Fig. 9A). However, the promotion of *GmNiR* expression was only significantly up-regulated by NaHS treatment from 5 to 28 DPI (RM GLM, *F* = 10.391, *P* = 0.029) (Fig. 9C). Moreover, NaHS treatment also up-regulated the expression levels of *GmSAT1* and *nifH* (RM GLM, *F* = 4.027, *P* = 0.041; *F* = 3.913, *P* = 0.046) (Fig. 9D, F). Though the expression levels of *GmAS* and *GmLb* did not show statistically significant variation during the entire treatment, they were slightly up-regulated by NaHS treatment at most of the checkpoints in the inoculated roots (RM GLM, *F* = 3.301, *P* = 0.192, *F* = 2.782, *P* = 0.210) (Fig. 9B, E).

**Figure 9.**
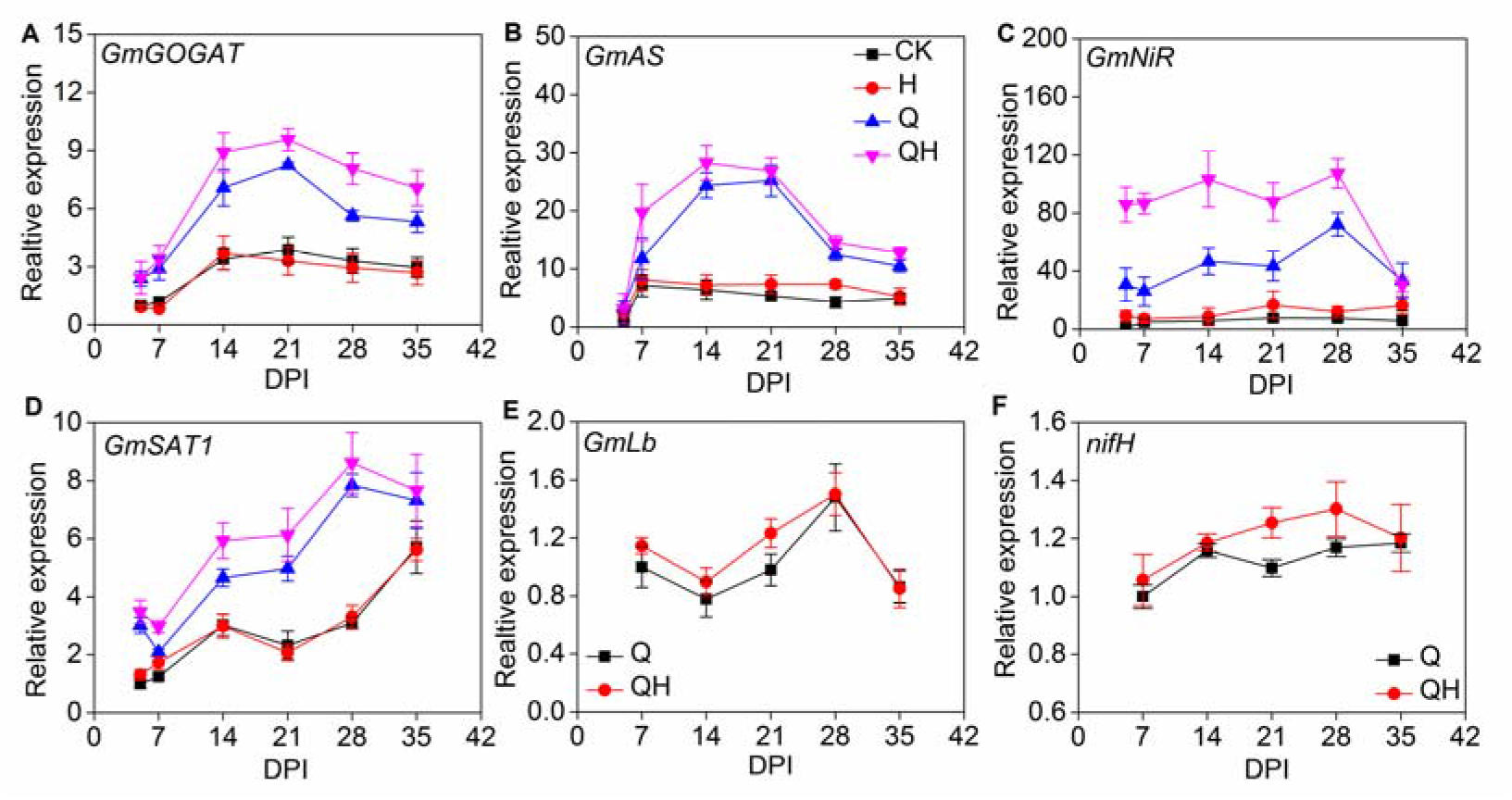
Gene expression level of nitrogen metabolism related genes. Relative expression levels of *GmGOGAT* (A), *GmAS* (B), *GmNiR*(C), and *GmSAT1*(D), *GmLb* (E) and *nifH* (F) are displayed in multiple-line charts with symbols. CK, controls. H, soybean plants treated with 100 μM NaHS. Q, Soybean plants inoculated with the *Sinorhizobium fredii* Q8 strain. QH, Soybean plants inoculated with the *Sinorhizobium fredii* Q8 strain and treated with 100 μM NaHS. The expression of *GmGOGAT, GmAS, GmNiR* and *GmSAT1* genes were relative to the expression level of respective gene in CK roots at 0.5 DPI. The expression of *GmLb* and *nifH* were relative to the expression level of respective gene in Q nodules at 7 DPI. Values are means ± SE (n = 9).

## Discussion

### H_2_S promotes soybean plant growth in the soybean-rhizobia symbiotic system

NaHS has been used as an exogenous H_2_S donor to study the physiological effects of H_2_S in many studies (Tamizhselvi et al., 2007; Wang et al., 2010; Li et al., 2012; Shi et al., 2013). To find the optimal concentration of NaHS for the growth of soybean, different concentrations (0, 10, 25, 50, 100, 250 and 500 μM) were used to treat soybean plants and *S. fredii* in liquid culture medium in our preliminary experiments. For *S. fredii*, 100 μM NaHS was shown to be the maximal concentration that did not significantly impact rhizobial growth (Fig. S1). For soybean plants, 100 μM NaHS was the optimal concentration for root length, shoot length, and biomass yield (Fig. S2). These preliminary results were in agreement with those of previous studies by Chen et al. (2011), Christou et al. (2014), and Tamizhselvi et al. (2007) using other plant materials. The ability of releasing H_2_S and increase the endogenous H_2_S concentration in soybean tissues (roots and shoots) by 100 μM NaHS was confirmed using the specific SF7-AM probe (Fig. 3 and Fig. S3).

Numerous experiments have shown that H_2_S can cause various biological effects in plants. In addition to alleviation of abiotic stresses such as heavy metals, heat, drought, and mineral salts (Zhang et al., 2008; Li et al., 2012; Shi et al., 2013; Chen et al., 2016), H_2_S was also found to be involved in regulating plant growth via different mechanisms (Chen et al., 2011; Zhang et al., 2011; Dooley et al., 2013). In the present study, we found that 100 μM NaHS promoted the growth of soybean plants under symbiotic conditions with the *S. fredii*(Fig. 1). In the inoculated soybean plants, NaHS treatment increased the root length and dry weight of roots (Fig. 1A, C). This indicated that H_2_S promoted the formation and development of roots, which is consistent with the findings of Zhang et al. (2009) in sweet potato. However, the length and biomass of shoots was not significantly affected by NaHS treatment. (Fig. 1B, D). As the total sulfur content in NaHS treatment is much lower than that in nitrogen-free nutrient solution (1:43), we may rule out the possibility that such changes in soybean growth were caused by extra addition of sulfur. Besides, Kalloniati et al. (2015) suggested that N-fixing nodules may act as source of reduced S and enhance the S assimilation of the legume plants. Together, these results indicated a potential role for H_2_S in regulating the symbiosis between *G. max* and *S. fredii*, which could promote symbiotic nitrogen fixation and consequently plant growth of soybean.

### H_2_S promtoes the formation of root nodules and nase activity

Nitrogen-fixing nodules formed by the symbiosis of rhizobia and legume plants are essential for fixation of nitrogen in the environment (Lee et al., 2014). The nodulation process is regulated by two critical systemic signaling events. The first is the recognition of the two symbiotic partners and the initiation of nodulation, which occurs in the rhizosphere. The second occurs within the plant to regulate the number of nodules as a means of balancing the resource cost and nitrogen benefit associated with nodulation (Hayashi et al., 2013). Many small molecules such as NO, H_2_O_2_, and phytohormones have been reported to be involved in the regulation of nodulation between legumes and rhizobia (Hirsch et al., 1997; Hérouart et al., 2002; Puppo et al., 2013). Here, we demonstrated that exogenous H_2_S increased the average number of nodules and the Nase activity in soybean roots (Fig. 2A, B).

In the *M. truncatula*-*S. meliloti* system, Del Giudice et al. (2011) confirmed the crucial position of NO in successful infection. Studies in *Arabidopsis* have shown that auxin accumulation is a prerequisite for organ formation in plants (Ferguson & Mathesius, 2014). External treatment of roots with auxins and auxin action inhibitors suppressed nodule formation, indicating that auxins are required for nodulation within a certain concentration range in *Medicago truncatula* (van Noorden et al., 2006). Infection event assays showed that during the early stages of the establishment of symbiosis, a larger number of infection events occurred in NaHS-treated soybean roots (Fig. 2C, D), suggesting that H_2_S may regulate the bacterial infection in the nodulation process, and may enhance the colonization of *S. fredii* into soybean nodules. Nase is the essential enzyme for symbiotic nitrogen fixation that catalyzes the conversion of dinitrogen into ammonia (Fishbeck et al., 1973). In the present study, Nase activity in NaHS-treated nodules was enhanced (Fig. 2B). The Nase complex consists of two metalloproteins that are highly conserved in sequence and structure throughout nitrogen-fixing bacteria (Halbleib & Ludden, 2000). One protein component contains the active site for substrate reduction with a molybdenum-iron (MoFe) cofactor, while the other iron (Fe)-containing protein acts as election donor to the MoFe component (Roth et al., 2010). The Fe protein is a 64 kDa α_2_ dimer of the *nifH* gene product (Halbleib & Ludden, 2000). In this research, we detected the nifH protein as an indicator for Nase content using western blotting analysis. The results showed that the abundance of the nifH protein was significantly increased by NaHS treatment (Fig. 7A, C). And this result is in coincidence with the result by qRT-PCR analysis of *nifH* gene expression (Fig. 9F). NaHS treatment increased the transcription abundance of *nifH* during the treatment (Fig. 9F). Together, the results of western blot analysis and qRT-PCR indicated that the NaHS treatment promoted the Nase activity through up-regulating the expression of *nifH* gene and increasing Fe protein content in soybean root nodules.

Baudouin et al. (2006) demonstrated that NO is synthesized in *M. truncatula* nodules, and is required for nodule functioning, and Leach et al. (2010) used NO synthase-specific inhibitors to show that NO is crucial for the development of functional nodules in soybean. In our current study, transmission electron microscopy was conducted to observe the microscopic structure of infected cells, symbionts, and bacteroids. Interestingly, bacteroids without a surrounding peribacteroid membrane were present in the intercellular space at 14 DPI in NaHS-treated nodules (Fig. 4B). It was worth noting that no typical symbiont structure was formed by these intercellular bacteroids. Thus, H_2_S may act as a signaling molecule in regulating the localization and differentiation of bacteroids in root nodules. Collectively, our results suggested H_2_S was closely related to rhizobial infection, bacteroid differentiation, nitrogen fixation, and nodule development in the soybean-rhizobia symbiotic system.

### H_2_S promotes photosynthetic and photochemical activity in soybean plants

Herein, we demonstrated the regulatory role of H_2_S in symbiosis, nitrogen fixation, and metabolism in the soybean-rhizobia symbiotic system. In plants, adequate nitrogen content can positively affect many physiological processes, such as flowering, carbon assimilation in tissues, and ion uptake (Fernandes & Rossiello, 1995; Xu et al., 2001; Liu et al., 2008). In addition, nitrogen content is closely related to chlorophyll content and photosynthesis in plant leaves (Gulmon & Chu, 1981; Lapointe, 1987), because nitrogen is a composition of chlorophyll and photosynthesis-related proteins. Thereby, nitrogen content in soybean plants may influence the formation of chloroplasts and accumulation of chlorophyll in them. In this work, both chlorophyll content and photosynthetic rate were elevated in leaves of NaHS-treated plants (Fig. 5). Additionally, soybean plants were cultured in perlite-vermiculite substrate and watered with nitrogen-free nutrient solution. It caused nitrogen starvation during plant growth. As H_2_S may promote soybean nodulation and nitrogen fixation ability, it may increase the nitrogen supplement in the soybean plants, which eventually alleviated nitrogen starvation and increased chlorophyll content and photosynthetic rate in the NaHS treated soybean plants. Although, H_2_S significantly increased the chlorophyll content in the soybean leaves, it did not significantly affect the NH_4_^+^ and NO_3_^−^ concentration in soybean roots and shoots (Fig. S6). This result is understandable, that under nitrogen deficiency conditions, limited nitrogen in plant tissues will be rapidly transformed into organic compounds.

H_2_S also acted as a modulator of PSII activity in soybean. Specifically, PSII, Fv/Fm, and Fv′/Fm′ parameters in inoculated plants were increased (Fig. 6A, C, and D), indicating that the photochemical efficiency of PSII was increased. These are consistent with the increase in chlorophyll content caused by 100 μM NaHS treatment. Together with increased chlorophyll content and PSII activity, this enhanced photosynthesis led to higher carbon assimilation, and ultimately to increased biomass of soybean.

### H_2_S regulates the expression of genes and proteins related to symbiosis and nitrogen metabolism

After analysis at the phenotypic level, we sought to elucidate the effects of H_2_S on the expression of genes and proteins related to symbiosis and nitrogen fixation in the soybean-rhizobia system. Firstly, we focused our attention on CHS, an enzyme catalyzing the synthesis of chalcone, which is reported to be a key precursor of a series of flavonoids (Schijlen et al., 2007; Dao et al., 2011). Li et al. (1993) demonstrated that CHS-deficient *Arabidopsis* displayed lower production of flavonoids. In the present study, the results of western blotting analysis showed that the abundances of CHS in root nodules were not affected by H_2_S (Fig. 7A, B).

Then, we investigated the expression of genes related to symbiosis and nitrogen metabolism in root nodules. *GmENOD40* was reported to be the downstream components of the perception of NFs (Ferguson et al., 2010), which is expressed in pericycle cells of root vascular bundles, dividing cortical cells, the nodule primordium, and developing nodules (Ferguson & Mathesius, 2014). Charon et al. (1999) reported that alteration in the expression of *ENOD40* could influence the nodulation, suggesting that it plays a vital role in in nodule organogenesis. In the present study, expression of *GmENOD40* gene in soybean was up-regulated by NaHS treatment (Fig. 8A). *NIN* is essential for nodule organogenesis and is also required for the initiation of bacterial infection in the roots (Madsen & Al, 2009; Vernié et al., 2015). In this case, the expression levels of three *NIN* genes in soybean were up-regulated by NaHS treatment (Fig. 8D, E, F). Besides, the other two nodulation marker genes involved in the NFs nodulation pathway, *GmERN* and *GmNSP2b* were also activated in the NaHS treated soybean roots (Fig. 8B, C). Together, the present results suggested that H_2_S stimulated the expression of *GmENOD40, GmERN, GmNSP2b*, and *GmNIN* genes. As these genes play a crucial role in the NFs nodulation signaling pathway and are closely related to the organogenesis and development of root nodules. These results may elucidate the underlying mechanisms of enhanced nodulation and nodule structural changes, that NaHS promoted soybean nodulation and nodule development through regulating the expression abundances of symbiotic related genes.

On the bacterial side, *Nod* genes in rhizobia are crucial for the early stages of recognition by legume hosts because they encode host-specific lipochito-oligosaccharidic Nod factors (NF) that can activate downstream symbiotic reactions by binding to plant kinase-like receptors (Giraud et al., 2007). Among the *nod* genes, *nodD* is a key player, because after sensing flavonoid signals secreted by legume roots into the soil, *nodD* initiates the expression of other *nod* genes (Machado & Krishnan, 2003). The results of qRT-PCR demonstrated that transcript abundances of *nodC* and *nodD* genes were increased in NaHS-treated nodules, suggesting H_2_S does not only promote symbiosis by affecting the legume host, but also triggers symbiotic responses in the rhizobia.

Uptake of nitrate by root cells followed by reduction and assimilation in plant tissues is the main route by which mineral nitrogen is converted into organic nitrogen by living organisms. Like photosynthesis, these are life-dependent processes (Imsande & Touraine, 1994). In our study, the relative expression levels of six genes related to nitrogen metabolism, including soybean genes *GmGOGAT, GmAS, GmNiR, GmSAT1, GmLb*, and rhizobial gene *nifH*, were quantified by qRT-PCR. These genes encode crucial players in amino acid metabolism and nitrogen assimilation. Expression of *GmGOGAT, GmAS, GmNiR*, and *GmSAT1* was strongly induced by H_2_S compared to untreated controls. Glutamic acid and aspartic acid are two essential amino acids that provide carbon skeletons for the synthesis of many amino acids by transamination. Indeed, δ-aminolevulinic acid, a precursor of chlorophyll, is synthesized from the intact carbon skeleton of glutamic acid (Beale, 1990). Furthermore, asparagine is the common precursor of the essential amino acids lysine, threonine, methionine, and isoleucine in higher plants, and aspartate may also be converted to asparagine in a potentially competing reaction (Azevedo et al., 2006). The ammonium transporter encoded by *GmSAT1* was reported to be located in the peribacteroid membrane, and is believed to be responsible for the transportation of ammonium fixed by bacteroids in soybean nodules (Kaiser et al., 1998). Thus, up-regulation of *GmGOGAT, GmAS*, and *GmSAT1* indicates that H_2_S might enhance the transportation of fixed ammonium and its assimilation into amino acids. In addition, nitrate assimilation can occur in either the roots or leaves via the nitrate reductase-nitrite reductase pathway (Evans, 2001). The product of this assimilation event is NH_4_^+^, and assimilation of NH_4_^+^ occurs through the glutamine synthetase-glutamate synthase pathway. Assimilation occurs in the roots, near the site of uptake, to avoid toxic accumulation (Bloom, 1988). Therefore, we can conclude that Therefore, we can conclude that H_2_S up-regulated the expression of *GmGOGAT, GmAS, GmNiR* and *GmSAT1 genes*, through which led to higher N assimilation.

## Conclusion

In this study, we demonstrated the potential role of H_2_S in symbiosis and nitrogen fixation in the *G. max*-*S. fredii* symbiotic system. We hypothesized a model to explain the stimulatory effects of H_2_S on symbiosis. As shown in Figure 10, our results suggested that H_2_S enhances the symbiotic relationship of *G. max*-*S. fredii* as NaHS treatment promoted nodulation and Nase activity in soybean. Besides, H_2_S promoted infection of rhizobia into both roots and nodules of soybean by up-regulating the expression of symbiosis related genes, such as *GmENOD40, GmERN, GmNSP2b, GmNIN* genes, and *nodC*. Additionally, the abundance of nifH protein was increased in soybean nodules following NaHS treatment. Moreover, NaHS treatment stimulated the expression of N metabolism related genes, such as *GmGOGAT, GmNR, GmSAT1* and *nifH*. The enhanced nitrogen fixation and assimilation increased the chlorophyll content, the photosynthetic rate, and PSII activity in inoculated plants. Finally, this enhanced photosynthesis endowed soybean plants with greater biomass and more vigorous growth. Taken together, these findings suggested H_2_S promoted soybean growth under symbiotic conditions by enhancing the establishment of symbiosis and stimulating nitrogen fixation in the *G. max*-*S*. *fredii* symbiotic system. Further details of the underlying molecular mechanisms, such as the targets of H_2_S, and how H_2_S fits into regulatory signaling pathways, still need further study.

**Figure 10.**
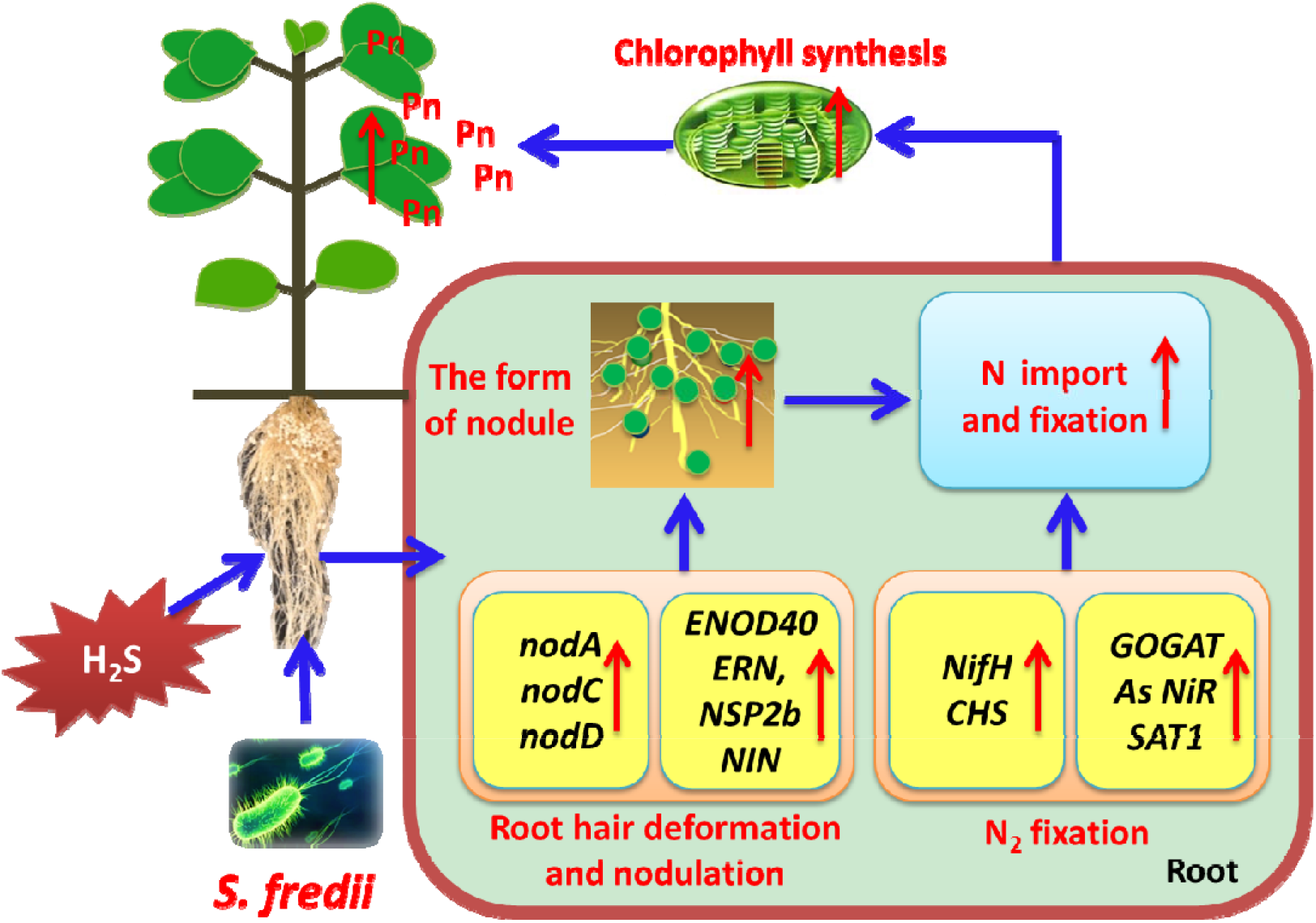
Schematic model of the mechanisms underlying the positive effects of H_2_S on the *Glycine max*-*Sinorhizobium fredii* symbiotic system. The model, based on the results of the present study and the wider literature, explains how exogenous H_2_S might influence the nodulation signaling pathway and the biological nitrogen fixation capacity in *G. max*.

## Supplemental Data

The following materials are available on the online version of this article.

**Table S1.** Nucleotide sequences of primers used in the qRT-PCR assay.

**Table S2.** qRT-PCR programs used for relative expression assay of target genes.

**Figure S1.** Effect of different concentrations of NaHS treatment on cell growth of *Sinorhizobium fredii*.

**Figure S2.** Effect of different concentrations of NaHS treatment on plant growth of *Glycine max*.

**Figure S3.** Quantification of fluorescence intensity of SF7-AM in soybean tissues with and without NaHS treatment.

**Figure S4.** Light microscopy analysis of infection events in soybean roots.

**Figure S5** Light microscopy analysis of paraffin sections of root nodules.

**Figure S6** H_2_S’s effect on the concentration of NH_4_^+^ and NO_3_^−^ in soybean roots and Shoots.

## Acknowledgements

We are grateful to Yan-Tao Luo, Ming-Mei Lu and Jian-Qiang Liang for guidance on experimental methods. Shuo Jiao has provided constructive suggestions to the writing of this article. This study was financially supported by the Natural Science Foundation of China (NSFC) (31501822) and the Postdoctoral Science Foundation of China (2015M580876 and 2016T90948).

## Abbreviations

Ci: carbon dioxide concentration.
Fm: maximal fluorescence yield.
Fo: minimal fluorescence yield.
Gs: stomatal conductance.
Pn: photosynthetic rate.
PSII: photosystem II.
qRT-PCR: quantitative real-time PCR.
Tr: transpiration rate.
Vpdl: vapor pressure divicit.

## References

Abdel-Lateif K, Vaissayre V, Gherbi H, Verries C, Meudec E, Perrine-Walker F, Cheynier V, Svistoonoff S, Franche C, Bogusz D. 2013. Silencing of the chalcone synthase gene in *Casuarina glauca* highlights the important role of flavonoids during nodulation. New Phytologist 199, 1012–1021

Álvarez C, García I, Moreno I, Pérez-Pérez ME, Crespo JL, Romero LC, Gotor C. 2012. Cysteine-generated sulfide in the cytosol negatively regulates autophagy and modulates the transcriptional profile in *Arabidopsis*. Plant Cell 24, 4621–4634

Azevedo RA, Lancien M, Lea PJ. 2006. The aspartic acid metabolic pathway, an exciting and essential pathway in plants. Amino Acids 30, 143–162

Baudouin E, Pieuchot L, Engler G, Pauly N, Puppo A. 2006. Nitric oxide is formed in *Medicago truncatula-Sinorhizobium meliloti* functional nodules. Molecular Plant-Microbe Interactions 19, 970–975

Beale SI. 1990. Biosynthesis of the tetrapyrrole pigment precursor, δ-aminolevulinic acid, from glutamate. Plant Physiology 93, 1273–1279

Becana M, Aparicio-Tejo PM, Sánchez-Díaz M. 1985. Nitrate and nitrite reduction by alfalfa root nodules: Accumulation of nitrite in *Rhizobium melioti* bacteroids and senescence of nodules. Physiologia Plantarum 64, 353–358

Becana M, Sprent JI. 1987. Nitrogen fixation and nitrate reduction in the root nodules of legumes. Physiologia Plantarum 70, 757–765

Benyamina SM, Baldaccicresp F, Couturier J, Chibani K, Hopkins J, Bekki A, De LP, Rouhier N, Jacquot JP, Alloing G. 2013. Two *Sinorhizobium meliloti* glutaredoxins regulate iron metabolism and symbiotic bacteroid differentiation. Environmental Microbiology 15, 795

Bloom AJ. 1988. Ammonium and nitrate as nitrogen sources for plant growth, Isi Atlas of Science-Animal & Plant Sciences 1, 55–59

Bradford MM. 1976. A rapid and sensitive method for the quantitation of microgram quantities of protein utilizing the principle of protein-dye binding. Analytical Biochemistry 72, 248–254

Caballero-Mellado J, Martinez-Romero E. 1999. Soil fertilization limits the genetic diversity of Rhizobium in bean nodules. Symbiosis 26 (2), 111–121

Cam Y, Pierre O, Boncompagni E, Hérouart D, Meilhoc E, Bruand C. 2012. Nitric oxide (NO): a key player in the senescence of *Medicago truncatula* root nodules. New Phytologist 196, 548–560

Charon C, Sousa C, Crespi M, Kondorosi A. 1999. Alteration of enod40 expression modifies *Medicago truncatula* root nodule development induced by *sinorhizobium meliloti*. Plant Cell 11, 1953–1965

Chen J, Shang YT, Wang WH, Chen XY, He EM, Zheng HL, Shangguan ZP. 2016. Hydrogen sulfide-induced drought tolerance in *Spinacia oleracea* seedlings. Frontiers in Plant Science 7,27–45

Chen J, Shang YT, Zhang NN, Zhong Y, Wang WH, Zhang JH, Shangguan Z. 2018. Sodium hydrosulfide modifies the nutrient ratios of soybean (*Glycine max*) under iron deficiency. Journal of Plant Nutrition and Soil Science 181, 305–315

Chen J, Wang WH, Wu FH, You CY, Liu TW, Dong XJ, He JX, Zheng HL. 2013. Hydrogen sulfide alleviates aluminum toxicity in barley seedlings. Plant and Soil 362, 301–318

Chen J, Wu FH, Wang WH, Zheng CJ, Lin GH, Dong XJ, He JX, Pei ZM, Zheng HL. 2011. ydrogen sulphide enhances photosynthesis through promoting chloroplast biogenesis, photosynthetic enzyme expression, and thiol redox modification in *Spinacia oleracea* seedlings. Journal of Experimental Botany 62, 4481–4493

Christou A, Filippou P, Manganaris GA, Fotopoulos V. 2014. Sodium hydrosulfide induces systemic thermotolerance to strawberry plants through transcriptional regulation of heat shock proteins and aquaporin. BMC Plant Biology 14, 42–42

Christou A, Manganaris GA, Papadopoulos I, Fotopoulos V. 2013. Hydrogen sulfide induces systemic tolerance to salinity and non-ionic osmotic stress in strawberry plants through modification of reactive species biosynthesis and transcriptional regulation of multiple defence pathways. Journal of Experimental Botany 64, 1953–1966

Damiani I, Pauly N, Puppo A, Brouquisse R, Boscari A. 2016. Reactive oxygen species and nitric oxide control early steps of the legume-rhizobium symbiotic interaction. Frontiers in Plant Science 7, 218–227

Dao T, Linthorst H, Verpoorte R. 2011. Chalcone synthase and its functions in plant resistance. Phytochemistry Reviews 10, 397

De BC, Mylona P, Yang WC, Katinakis P, Bisseling T, Franssen H. 1993. Characterization of the soybean early nodulin cDNA clone GmENOD55. Plant Molecular Biology 22, 1167–1171

Del Giudice J, Cam Y, Damiani I, Fung-Chat F, Meilhoc E, Bruand C, Brouquisse R, Puppo A, Boscari A. 2011. Nitric oxide is required for an optimal establishment of the *Medicago truncatula-Sinorhizobium meliloti* symbiosis. New Phytologist 191, 405–417

Dooley FD, Nair SP, Ward PD. 2013. Increased growth and germination success in plants following hydrogen sulfide administration. PLoS ONE 31, S24–S24

Evans RD. 2011. Physiological mechanisms influencing plant nitrogen isotope composition. Trends in Plant Science 6, 121

Fang HH, Liu ZQ, Jin ZP, Zhang LP, Liu DM, Pei YX. 2016. An emphasis of hydrogen sulfide-cysteine cycle on enhancing the tolerance to chromium stress in *Arabidopsis*. Environmental Pollution 213, 870–877

Farnham MW, Griffith SM, Miller SS, Vance CP. 1990. Aspartate aminotransferase in alfalfa root nodules. III. Genotypic and tissue expression of aspartate aminotransferase in alfalfa and other species. Plant Physiology 94, 1634–1640

Ferguson BJ, Indrasumunar A, Hayashi S, Lin MH, Lin YH, Reid DE, Gresshoff PM. 2010. Molecular analysis of legume nodule development and autoregulation. Journal of Integrative Plant Biology 52, 61–76

Ferguson BJ, Mathesius U. 2014. Phytohormone regulation of legume-rhizobia interactions. Journal of Chemical Ecology 40, 770–790

Fernandes MS, Rossiello ROP. 1995. Mineral nitrogen in plant physiology and plant nutrition. Critical Reviews in Plant Sciences 14, 111–148

Fishbeck K, Evans HJ, Boersma LL. 1973. Measurement of nitrogenase activity of intact legume symbionts in situ using the acetylene reduction assay. Agron J 65, 429–433

Fukuto JM, Carrington SJ, Tantillo DJ, Harrison JG, Ignarro LJ, Freeman BA, Chen A, Wink DA. 2012. Small molecule signaling agents: the integrated chemistry and biochemistry of nitrogen oxides, oxides of carbon, dioxygen, hydrogen sulfide, and their derived species. Chemical Research in Toxicology 25, 769–793

García-Mata C, Lamattina L. 2010. Hydrogen sulphide, a novel gasotransmitter involved in guard cell signalling. New Phytologist 188, 977–984

Giraud E, Moulin L, Vallenet D, Barbe V, Cytryn E, Avarre J-C, Jaubert M, Simon D, Cartieaux F, Prin Y. 2007. Legumes symbioses: absence of *Nod* genes in photosynthetic bradyrhizobia. Science 316, 1307–1312

Gulmon SL, Chu CC. 1981. The effects of light and nitrogen on photosynthesis, leaf characteristics, and dry matter allocation in the chaparral shrub, *Diplacus aurantiacus*. Oecologia 49, 207–212

Hérouart D, Baudouin E, Frendo P, Harrison J, Santos R, Jamet A, Van de Sype G, Touati D, Puppo A. 2002. Reactive oxygen species, nitric oxide and glutathione: a key role in the establishment of the legume-rhizobium symbiosis? Plant Physiology and Biochemistry 40, 619–624

Halbleib CM, Ludden PW. 2000. Regulation of biological nitrogen fixation. Journal of Nutrition 130, 1081–1084

Hichri I, Boscari A, Meilhoc E, Catalá M, Barreno E, Bruand C, Lanfranco L, Brouquisse R. 2016. Nitric oxide: A multitask player in plant-microorganism symbioses. In: L Lamattina, C García-Mata, eds. Gasotransmitters in Plants: The Rise of a New Paradigm in Cell Signaling. Springer International Publishing, Cham, pp 239–268

Hirsch A, Fang Y, Asad S, Kapulnik Y. 1997. The role of phytohormones in plant-microbe symbioses. Plant and Soil 194, 171–184

Imsande J, Touraine B. 1994. N demand and the regulation of nitrate uptake. Plant Physiology 105, 3

Jain M, Nijhawan A, Tyagi AK, Khurana JP. 2006. Validation of housekeeping genes as internal control for studying gene expression in rice by quantitative real-time PCR. Biochemical and Biophysical Research Communications 345, 646–651

Jamet A, Mandon K, Puppo A, Hérouart D. 2007. H_2_O_2_ is required for optimal establishment of the *Medicago sativa/Sinorhizobium meliloti* symbiosis. Journal of Bacteriology 189, 8741–8745

Kaiser BN, Finnegan PM, Tyerman SD, Whitehead LF, Bergersen FJ, Day DA, Udvardi MK. 1998. Characterization of an ammonium transport protein from the peribacteroid membrane of soybean nodules. Science 281, 1202–1206

Kalloniati C, Krompas P, Karalias G, Udvardi MK, Rennenberg H, Herschbach C, Flemetakis E. 2015. Nitrogen-Fixing Nodules Are an Important Source of Reduced Sulfur, Which Triggers Global Changes in Sulfur Metabolism in Lotus japonicus. Plant Cell 27, 2384–2400

Kanayama Y, Watanabe I, Yamamoto Y. 1990. Inhibition of nitrogen fixation in soybean plants supplied with nitrate I. Nitrite accumulation and formation of nitrosylleghemoglobin in nodules. Plant and Cell Physiology 31, 341–346

Kato K, Kanahama K, Kanayama Y. 2010. Involvement of nitric oxide in the inhibition of nitrogenase activity by nitrate in *Lotus* root nodules. Journal of Plant Physiology 167, 238–241

Kouchi H, Hata S. 1993. Isolation and characterization of novel nodulin cDNAs representing genes expressed at early stages of soybean nodule development. Molecular Genetics and Genomics 238, 106

Krause, Weis E. 2003. Chlorophyll fluorescence and photosynthesis: The basics. Annual Review of Plant Physiology 42, 313–349

Lapointe BE. 1987. Phosphorus-and nitrogen-limited photosynthesis and growth of *Gracilaria tikvahiae* (Rhodophyceae) in the Florida Keys: an experimental field study. Marine Biology 93, 561–568

Laureano-Marín AM, Moreno I, Romero LC, Gotor C. 2016. Negative regulation of autophagy by sulfide in *Arabidopsis thaliana* is independent of reactive oxygen species. Plant Physiology 171 (2), 1378

Leach J, Keyster M, Du Plessis M, Ludidi N. 2016. Nitric oxide synthase activity is required for development of functional nodules in soybean. Journal of Plant Physiology 167, 1584–1591

Lee SG, Krishnan HB, Jez JM. 2014. Structural basis for regulation of rhizobial nodulation and symbiosis gene expression by the regulatory protein NolR. Proceedings of the National Academy of Sciences 111, 6509

Li J, Oulee TM, Raba R, Amundson RG, Last RL. 1993. *Arabidopsis* flavonoid mutants are hypersensitive to UV-B irradiation. Plant Cell 5, 171

Li L, Rose P, Moore PK. 2011. Hydrogen sulfide and cell signaling. Annual Review of Pharmacology and Toxicology 51, 169–187

Li L, Sinkko H, Montonen L, Wei G, Lindström K, Räsänen LA. 2012. Biogeography of symbiotic and other endophytic bacteria isolated from medicinal *G lycyrrhiza* species in China. FEMS Microbiology Ecology 79, 46–68

Li L, Wang Y, Shen W. 2012. Roles of hydrogen sulfide and nitric oxide in the alleviation of cadmium-induced oxidative damage in alfalfa seedling roots. BioMetals 25, 617–631

Li ZG, Gong M, Xie H, Yang L, Li J. 2012. Hydrogen sulfide donor sodium hydrosulfide-induced heat tolerance in tobacco (*Nicotiana tabacum L*) suspension cultured cells and involvement of Ca^2+^ and calmodulin. Plant Science 185–186, 185–189

Li ZG, Yang SZ, Long WB, Yamg GX, Shen ZZ. 2013. Hydrogen sulphide may be a novel downstream signal molecule in nitric oxide-induced heat tolerance of maize (*Zea mays L*.) seedlings. Plant, Cell and Environment 36, 1564–1572

Lin VS, Lippert AR, Chang CJ. 2013. Cell-trappable fluorescent probes for endogenous hydrogen sulfide signaling and imaging H_2_O_2_-dependent H_2_S production. Proceedings of the National Academy of Sciences 110, 7131

Lin Y-T, Li M-Y, Cui W-T, Lu W, Shen WB. 2012. Haem Oxygenase-1 is Involved in Hydrogen Sulfide-induced Cucumber Adventitious Root Formation. Journal of Plant Growth Regulation 31, 519–528

Machado D, Krishnan HB. 2003. *nodD* Alleles of *Sinorhizobium fredii* USDA191 differentially influence soybean nodulation, *nodC* expression, and production of exopolysaccharides. Current Microbiology 47, 0134–0137

Madsen LH, Al E. 2009. The molecular network governing nodule organogenesis and infection in the model legume *Lotus japonicus*. Nature Communications 1, 10

Masclaux-Daubresse C, Daniel-Vedele FO, Dechorgnat J, Chardon F, Gaufichon L, Suzuki A. 2010. Nitrogen uptake, assimilation and remobilization in plants: challenges for sustainable and productive agriculture. Annals of Botany 105, 1141–1157

Masson-Boivin C, Giraud E, Perret X, Batut J. 2009. Establishing nitrogen-fixing symbiosis with legumes: how many rhizobium recipes? Trends in Microbiology 17, 458–466

Meilhoc E, Cam Y, Skapski A, Bruand C. 2010. The response to nitric oxide of the nitrogen-fixing symbiont *Sinorhizobium meliloti*. Molecular Plant-Microbe Interactions 23, 748–759

Mergaert P, Uchiumi T, Alunni B, Evanno G, Cheron A, Catrice O, Mausset A-E, Barloy-Hubler F, Galibert F, Kondorosi A. 2006. Eukaryotic control on bacterial cell cycle and differentiation in the Rhizobium-legume symbiosis. Proceedings of the National Academy of Sciences 103, 5230–5235

Minami E, Kouchi H, Carlson RW, Cohn JR, Kolli VK, Day RB, Ogawa T, Stacey G. 1996. Cooperative action of lipo-chitin nodulation signals on the induction of the early nodulin, ENOD2, in soybean roots. Molecular Plant-Microbe Interactions 9, 574

Moran NA. 2006. Symbiosis. Current Biology 16, R866–R871

Nishikawa F, Endo T, Shimada T, Fujii H, Shimizu T, Omura M. 2008. Isolation and characterization of a *Citrus* FT/TFL1 homologue (*CuMFT1*), which shows quantitatively preferential expression in *Citrus* seeds. Journal of the Japanese Society for Horticultural Science 77, 38–46

Oldroyd GE. 2013. Speak, friend, and enter: signalling systems that promote beneficial symbiotic associations in plants. Nature Reviews Microbiology 11, 252–263

Papanatsiou M, Scuffi D, Blatt MR, García-Mata C. 2015. Hydrogen sulfide regulates inward-rectifying K^+^ channels in conjunction with stomatal closure. Plant Physiology 168, 29–35

Pii Y, Crimi M, Cremonese G, Spena A, Pandolfini T. 2007. Auxin and nitric oxide control indeterminate nodule formation. BMC Plant Biology 7, 21

Puppo A, Pauly N, Boscari A, Mandon K, Brouquisse R. 2013. Hydrogen peroxide and nitric oxide: Key regulators of the legume-Rhizobium and mycorrhizal symbioses. Antioxidants & Redox Signaling 18, 2202

Romero LC, Aroca MÁ, Laureano-Marín AM, Moreno I, García I, Gotor C. 2014. Cysteine and cysteine-related signaling pathways in *Arabidopsis thaliana*. Molecular Plant 7, 264–276

Roth LE, Nguyen JC, Tezcan FA. 2010. ATP-and iron-protein-independent activation of nitrogenase catalysis by light. Journal of the American Chemical Society 132, 13672–13674

Sasakura F, Uchiumi T, Shimoda Y, Suzuki A, Takenouchi K, Higashi S, Abe M. 2006. A class 1 hemoglobin gene from *Alnus firma* functions in symbiotic and nonsymbiotic tissues to detoxify nitric oxide. Molecular Plant-Microbe Interactions 19, 441–450

Schijlen EG, de Vos CR, Martens S, Jonker HH, Rosin FM, Molthoff JW, Tikunov YM, Angenent GC, van Tunen AJ, Bovy AG. 2007. RNA interference silencing of chalcone synthase, the first step in the flavonoid biosynthesis pathway, leads to parthenocarpic tomato fruits. Plant Physiology 144, 1520–1530

Scuffi D, Álvarez C, Laspina N, Gotor C, Lamattina L, García-Mata C. 2014. Hydrogen sulfide generated by L-cysteine desulfhydrase acts upstream of nitric oxide to modulate abscisic acid-dependent stomatal closure. Plant Physiology 166, 2065–2076

Shen J, Xing T, Yuan H, Liu Z, Jin Z, Zhang L, Pei Y. 2013. Hydrogen sulfide improves drought tolerance in *Arabidopsis thaliana* by microRNA expressions. PLoS ONE 8, e77047

Shi H, Ye T, Chan Z. 2013. Exogenous application of hydrogen sulfide donor sodium hydrosulfide enhanced multiple abiotic stress tolerance in bermudagrass (*Cynodon dactylon (L). Pers*.). Plant Physiology and Biochemistry 71, 226–234

Stougaard J. 2000. Regulators and regulation of legume root nodule development. Plant Physiology 124, 531

Sun J, Wang R, Zhang X, Yu Y, Zhao R, Li Z, Chen S. 2013. Hydrogen sulfide alleviates cadmium toxicity through regulations of cadmium transport across the plasma and vacuolar membranes in *Populus euphratica* cells. Plant Physiology Biochemistry 65, 67–74

Tamizhselvi R, Moore PK, Bhatia M. 2007. Hydrogen sulfide acts as a mediator of inflammation in acute pancreatitis: In vitro studies using isolated mouse pancreatic acinar cells. Journal of Cellular and Molecular Medicine 11, 315

Tian H, Lu C, Melillo J, Ren W, Huang Y, Xu X, Liu M, Zhang C, Chen G, Pan S. 2012. Food benefit and climate warming potential of nitrogen fertilizer uses in China. Environmental Research Letters 7, 044020

Tilman D, Cassman KG, Matson PA, Naylor R, Polasky S. 2002. Agricultural sustainability and intensive production practices. Nature 418, 671–677

Trinchant J-C, Rigaud J. 1982. Nitrite and nitric oxide as inhibitors of nitrogenase from soybean bacteroids. Applied and Environmental Microbiology 44, 1385–1388

Vernié T, Kim J, Frances L, Ding Y, Sun J, Guan D, Niebel A, Gifford ML, De C-NF, Oldroyd GE. 2015. The NIN Transcription Factor Coordinates Diverse Nodulation Programs in Different Tissues of the Medicago truncatula Root. Plant Cell 27, 3410–3424

Vincent R, Fraisier V, Chaillou S, Limami MA, Deleens E, Phillipson B, Douat C, Boutin J-P, Hirel B. 1997. Overexpression of a soybean gene encoding cytosolic glutamine synthetase in shoots of transgenic *Lotus corniculatus L*. plants triggers changes in ammonium assimilation and plant development. Planta 201, 424–433

Wang BL, Shi L, Li YX, Zhang WH. 2010. Boron toxicity is alleviated by hydrogen sulfide in cucumber (*Cucumis sativus L*.) seedlings. Planta 231, 1301–1309

Wang MJ, Cai WJ, Li N, Ding YJ, Chen Y, Zhu YC. 2010. The hydrogen sulfide donor NaHS promotes angiogenesis in a rat model of hind limb ischemia. Antioxidants & Redox Signaling 12, 1065–1077

Werner GD, Cornwell WK, Cornelissen JH, Kiers ET. 2015. Evolutionary signals of symbiotic persistence in the legume-rhizobia mutualism. Proceedings of the National Academy of Sciences 112, 10262–10269

Xu GH, Wolf S, Kafkafi U. 2001. Effect of varying nitrogen form and concentration during growing season on sweet pepper flowering and fruit yield. Journal of Plant Nutritions 24, 1099–1116

Yan Z, Hossain MS, Arikit S, Valdés-López O, Zhai J, Wang J, Libault M, Ji T, Qiu L, Meyers BC. 2015. Identification of microRNAs and their mRNA targets during soybean nodule development: Functional analysis of the role of *miR393j-3p* in soybean nodulation. New Phytologist 207, 748–759

Yuan Z, Zhang Z, Wang X, Li L, Cai K, Han H. 2017. Novel Impacts of functionalized multi-walled carbon nanotubes in plants: promotion of nodulation and nitrogenase activity in rhizobium-legume system. Nanoscale 9, 28–36

Zhang H, Hu LY, Hu KD, He YD, Wang SH, Luo JP. 2018. Hydrogen sulfide promotes wheat seed germination and alleviates oxidative damage against copper stress. Journal of Integrative Plant Biology 50, 1518–1529

Zhang H, Hu SL, Zhang ZJ, Hu LY, Jiang CX, Wei ZJ, Liu J, Wang HL, Jiang ST. 2011. Hydrogen sulfide acts as a regulator of flower senescence in plants. Postharvest Biology and Technology 60, 251–257

Zhang H, Jiao H, Jiang CX, Wang SH, Wei ZJ, Luo JP, Jones RL. 2010. Hydrogen sulfide protects soybean seedlings against drought-induced oxidative stress. Acta Physiologiae Plantarum 32, 849–857

Zhang H, Tang J, Liu XP, Wang Y, Yu W, Peng WY, Fang F, Ma DF, Wei ZJ, Hu LY. 2009. Hydrogen sulfide promotes root organogenesis in *Ipomoea batatas, Salix matsudana* and *Glycine max*. Journal of Integrative Plant Biology 51, 1086–1094

